# Conceptual and analytical approaches for modeling the developmental origins of inequality

**DOI:** 10.1101/2022.02.10.479998

**Authors:** Anup Malani, Elizabeth A. Archie, Stacy Rosenbaum

## Abstract

In many species, individuals who experience harsh conditions during development often have poor health and fitness outcomes in adulthood relative to peers who do not. There are two classes of evolutionary hypotheses for the origins of these early life contributors to inequality in adulthood: developmental constraints (DC) models, which focus on the deleterious effects of low-quality early-life environments, and predictive adaptive response (PAR) hypotheses, which emphasize the cost of mismatches between early and adult environments. Distinguishing DC and PAR models empirically is difficult for both conceptual and analytical reasons. Here, we resolve this difficulty by providing explicit mathematical definitions for DC, PARs, and related concepts, and propose a novel, quadratic regression-based statistical test derived from these definitions. Simulations show that this approach improves the ability to discriminate between DC and PAR hypotheses relative to a common alternative based on testing for interaction effects between developmental and adult environments. Simulated data indicate that the interaction effects approach often conflates PARs with DC, while the quadratic regression approach yields high sensitivity and specificity for detecting PARs. Our results highlight the value of linking verbal and visual models to a formal mathematical treatment for understanding the developmental origins of inequitable adult outcomes.

## 1 Introduction

Early life adversity is associated with a wide variety of negative outcomes in adulthood, including poor health, short lifespans, and low evolutionary fitness. These “early life effects” are believed to be a major contributor to inequalities in adult outcomes and thus have generated strong interest from many fields, including ecology and evolution [1, 2], public health and medicine [3, 4], psychology and psychiatry [5, 6], sociology [7], and economics [8]. The widespread observation of early life effects across species [9] has given rise to two broad classes of hypotheses that seek to explain the evolution of such effects [reviewed in 10, 11, 12]. The first class, often termed developmental constraints (DC) or “silver spoon” hypotheses [13, 14], posits that harsh conditions in early life decrease later-life phenotypic quality (and thus, evolutionary fitness), perhaps because organisms make tradeoffs in harsh developmental environments that promote immediate survival but compromise long-term outcomes [12]. The second class, predictive adaptive response (PAR) hypotheses, proposes that there is selection for organisms to use their early life conditions to predict characteristics of their adult environment and adjust their responses to the early environment accordingly [14, 15]. Under this model, incorrect “guesses” about the future lead to poor outcomes during adulthood. Organisms may predict their external environment (referred to as an external PAR), or the quality of their own somatic state (an internal PAR) [15, 16]).

A key difference between DC and PAR hypotheses is whether the early environment *per se* is the primary determinant of later life outcomes, or whether the *difference* (i.e., mismatch) between early and adult environments is the determining factor. DC hypotheses focus on the former (Figure 1a): poor-quality developmental environments are predicted to lead to worse outcomes in adulthood [14]. In contrast, PAR hypotheses focus on the effects of *change* in the environment on adult outcomes (Figure 1b), and more specifically, the costs of incorrectly predicting whether a change will occur. A shared feature of all PAR hypotheses is the idea that organisms use cues available during development to make predictions about some feature of adulthood (e.g., what their environment will be like, or what their somatic state will be). The less accurate the prediction, the worse an organism’s outcomes will be [17, 18].

**Figure 1:**
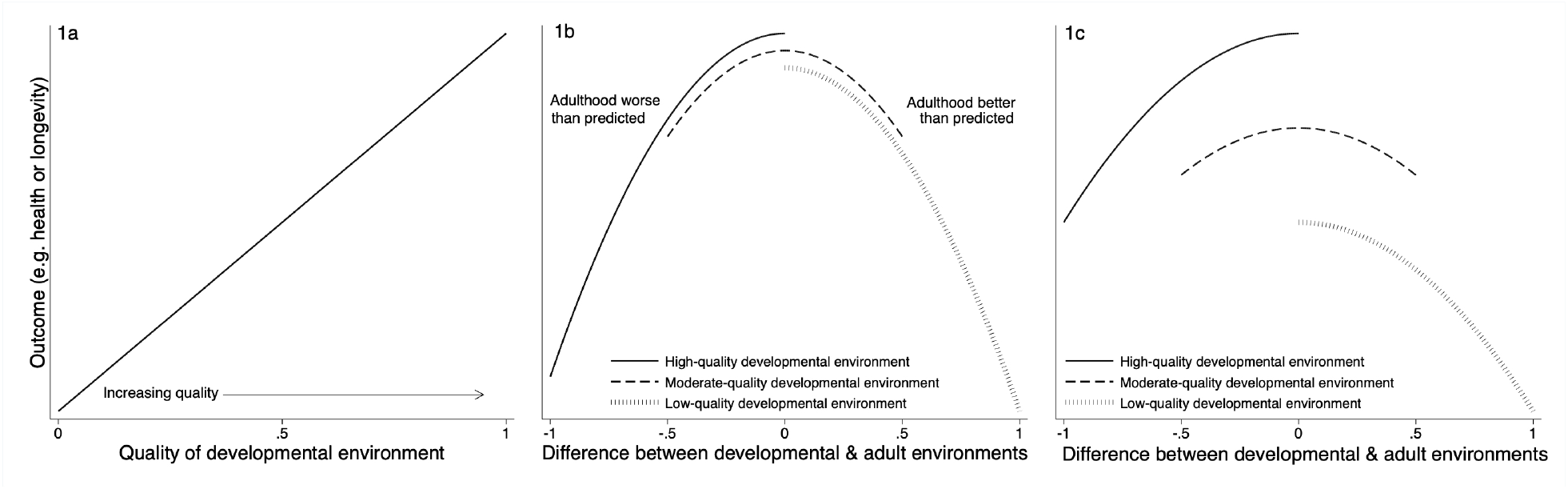
Visualizations of the developmental constraints and predictive adaptive response hypotheses If environmental quality is a continuous variable bounded by 0 (worst) and 1 (best), **(1a)** the developmental constraints (DC) hypothesis predicts that low-quality developmental environments will result in worse adult outcomes (e.g., health or longevity) than high-quality environments. **(1b)** The predictive adaptive response (PAR) hypothesis predicts that, controlling for adult environmental quality, bigger differences between an organism’s developmental and adult environments leads to worse adult outcomes. The center of the x-axis (0) represents a perfect match between developmental and adult environment. Negative x-axis values (bounded by −1) represent an organism who predicted a high-quality adult environment (based on their developmental environment) but ended up in a low-quality environment. Positive x-axis values (bounded by 1) represent an organism who predicted a low-quality adult environment but ended up in a high-quality environment. An organism that experienced a moderate-quality developmental environment can experience an adult environment is either worse or better than they predicted it would be, resulting in negative or positive x-axis values, respectively (bounded by −0.5 and 0.5). **(1c)** The DC and PAR hypotheses, as defined in Section 2, can be true simultaneously. An organism who experienced a low-quality developmental environment can have worse outcomes relative to what their outcomes would have been had they experienced a high-quality developmental environment, <u>and</u> can fare worse the greater the difference between their developmental and adult environments.

As conceptualized above, the DC and PAR hypotheses are not mutually exclusive, as many others have noted [19, 20, 11, 21]. Adult outcomes can simultaneously be determined by the quality of the developmental environmental *and* how well organisms predict some feature of adulthood (Figure 1c) [18, 22]. For example, depending on the size of the effects of developmental constraints, an organism that starts off in a low-quality environment may always fare worse than if it had started in a high-quality environment, even if it would <u>also</u> fare worse if there were a mismatch between its early and adult environments (Figure 1c). In parallel, higher-quality adult environments may also yield better outcomes independent of the developmental environment or an organism’s predictive accuracy [23, 20]. This conceptually related idea, termed the adult environment quality hypothesis (AEQ) [14], is implicit in tests of the relationship between adult environment and fitness that do not consider early life.

Currently, in humans and other long-lived mammals, DC hypotheses have stronger empirical support than PAR hypotheses [reviewed in 11]. However, it is difficult to know what conclusions to draw from this pattern because the ubiquitous verbal and visual models used for testing these hypotheses create uncertainty about precisely what is being evaluated. Explicit mathematical definitions for DC, PAR, and AEQ models can help remedy these problems and resolve ambiguity in interpreting the empirical data [24, 25]. Here, we provide such definitions and use them to motivate a quadratic regression model that tests these hypotheses. Using simulations, we demonstrate substantial improvements in the sensitivity and specificity of tests for DC and PAR with our quadratic regression approach over a commonly used interaction regression model [14, 26]. We show that a key reason for this improvement is that, unlike the interaction regression approach, the quadratic regression method avoids conflating DC with PARs. Additionally, we discuss conceptual issues that continue to constrain formal tests of DC and PAR models, and provide guidance on practical implementation of tests for these hypotheses.

## 2 Mathematical definitions and predictions

In this section we propose formal definitions for DC, PAR, and the related adult environmental quality (AEQ) hypothesis mentioned briefly above. We then outline the theoretical predictions that follow from these definitions, and formalize assumptions that are often left implicit in studies of the early life origins of inequality. Our definitions and predictions are summarized in Table 1.

**Table 1:** Formal definitions and derived predictions of theories for the relationship between the quality of developmental environments and inequalities in adult outcomes.

| Theory | Definition | Observable Variation | Prediction or Test (Eqn. No.) |
| --- | --- | --- | --- |
| Developmental Constraints (DC) | $\frac{\partial y_1}{\partial e_0} > 0$ | $(y_1, e_0)$ | $\frac{\partial y_1}{\partial e_0} > 0$ (1) |
| Predictive Adaptive Response (PAR) | $E(e_1) = e_0$<br>$\frac{\partial p}{\partial E(e_1)} < 0$<br>$\frac{\partial^2 y_1}{\partial p \partial e_1} < 0$ | | |
| Phenotypic adaptation test | | $(e_0, p)$ | $\frac{\partial p}{\partial e_0} < 0$ (5) |
| Environmental mismatch test | | $(y_1, \vert \Delta e\vert )$ | $\frac{\partial y_1}{\partial \vert \Delta e\vert } < 0$ (7) |
| Adult Environmental Quality (AEQ)* | $\frac{\partial y_1}{\partial e_1} > 0$ | $(y_1, e_1)$ | $\frac{\partial y_1}{\partial e_1} > 0$ (8) |

$y_0$ = developmental outcome; $y_1$ =outcome in adulthood; $e_0$ =developmental environment; $e_1$ =adult environment; $E(e_1)$ =adult environment the organism expects; $\Delta e$ =difference between developmental and adult environments; $p$ =phenotypic adaptation to a given environment. Definitions assume a differentiable function that maps from environment or discrepancy between environment and adapted-to environment to outcomes. See Section 2 for details.

We note three conceptual points about our definitions. First, they posit what would happen if, for a given organism, its developmental environment were to be different, or if it made a different prediction about its adult environment. A useful way to phrase the inquiry is: how would a given individual’s outcome differ if, in a parallel universe, its environment were different? Our definitions are not about the distributions of environments and outcomes across individuals in a given population. For example, we will not define DC as a claim that organisms with better outcomes tend to have been born in better environments.

These clarifications are required because, though our definitions are intra-individual (in the sense that they compare an individual to itself in another hypothetical world), the DC and PAR hypotheses are often empirically tested with data that relies on inter-individual comparisons. That is, *tests* of the theories we define often rely on data that compares one individual to another. Whether it is appropriate to use inter-individual data to test intra-individual hypotheses is a methodological question concerning causation that is beyond the scope of this paper.

Second, our definitions constitute theories or hypotheses about the influence of early environment on an organism’s environment, and they generate (testable) predictions about how environments affect outcomes. In some cases the theory and the prediction are the same: the prediction of the theory is mathematically identical to the statement of the theory. An example is DC, where the theory says low-quality early environments cause poor adult outcomes and the prediction is that low-quality early environments cause poor adult outcomes. In other cases, such as PAR, the definitions and the predictions are not the same.

Third, our definitions assume that developmental and adult environment are continuous variables and that one can take derivatives of the functions that relate those environments to outcomes. With some technical modifications, our conclusions can be extended to categorically-coded variables.

### Developmental constraints (DC) hypothesis

The DC hypothesis proposes that lower-quality developmental environments will lead to worse outcomes in adulthood, relative to higher-quality developmental environments [14]. For example, *Drosophila melanogaster* larvae which are nutritionally stressed become smaller adults and have lower egg viability in adulthood than larvae which do not [27].

DC can be described by a simple, within-individual causal chain:

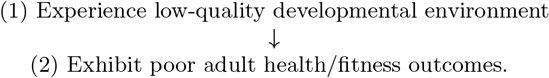

Mathematically, this can be represented as:

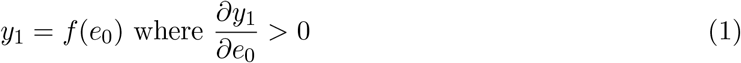

where *y*_1_ is adult outcomes (e.g., health or longevity), *f* is a decreasing function, and *e*_0_ is the developmental environment. Higher values of *y*_1_ and *e*_0_ indicate better outcomes or higher-quality environments, respectively. DC can be directly tested with empirical data, assuming that the developmental environment is independent from other factors that predict adult health and fitness. Our definition of DC is evolutionary in the sense that it proposes that early life adversity is a potential selective force, but it takes no stance on the mechanism(s) by which early adversity is connected to worse adult outcomes.

### 2.2 Predictive adaptive response (PAR) hypothesis

PAR hypotheses propose that an organism develops a phenotype optimized for its expected adult environment [28, 15]. Different versions of the PAR hypothesis distinguish between predictions based on cues in the external environment (e.g., temperature or rainfall, an external PAR) or based on internal somatic state (i.e., an internal PAR). Evidence that meadow voles who mature in colder environments are born with thicker coats has been widely interpreted as evidence of an external PAR, because the benefits of the thicker coat phenotype primarily accrue in adulthood [29]. A related hypothesis, the developmental mismatch hypothesis, is ubiquitous in the evolutionary health literature [16, 30].^1^ It posits that the more closely the environment at life-history stage *m* matches the environment to which the organism adapted at some prior stage *m − k* for *k >* 0, the better off an organism’s outcomes at stage *m* will be.

The within-individual causal chain that motivates the PAR hypothesis and may motivate the mismatch hypothesis is expressed as a set of four steps:

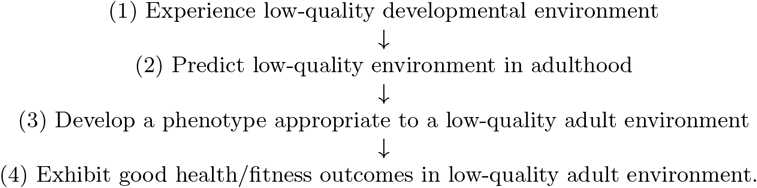

The above assumes that all else is held constant, and that there is an analogous causal chain for high-quality environments.

Each step in this chain is necessary. The proximate mechanism underlying PAR is captured by steps 2 to 3: the organism makes a prediction about its adult environment, and that prediction must lead it to adjust to cope with that predicted future environment. Step 4 is required to complete the evolutionary logic of PAR (i.e., the “ultimate” mechanism): this response evolves because adopting a phenotype optimized to a low-quality adult environment improves fitness outcomes for an organism in adulthood relative to the outcomes it would have experienced if it had not adopted that phenotype.

Steps 1 and 4 typically correspond to observables in real data sets. Step 1 can be captured by measures of environmental variation where some values are thought to be high-quality/benign and some low-quality/adverse. Step 4 can be captured by measures of health or fitness-related outcomes. Step 3 is theoretically observable, if the relevant, optimizable phenotypic response is known (e.g., smaller body size if the environment is food-restricted). However, step 2, an organism’s prediction, is not typically observable, so step 1 serves as a proxy for an organism’s prediction:

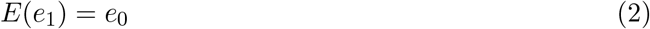

This says that an organism expects that its adult environment will be the same as its developmental environment. Equation 2 is not essential to defining PAR, but unlike an organism’s prediction, the developmental environment is observable, making the PAR hypothesis falsifiable.

The steps outlined above can be mapped to mathematical expressions. PAR is defined by two propositions. The first captures steps 2 to 3:

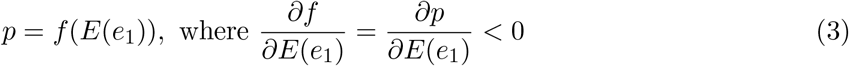

where *p* is the value of a given phenotype that is protective in low quality environments and *e*_1_ is the quality of the adult environment. (Our choice to define the adaptation to high *e*_1_ to be low *p*, i.e., to suppose that it is a bad environment that needs an adaptation, is arbitrary. We could have assumed that high *p* is better suited for high *e*_1_ so long as we also reversed the signs of *∂p/∂E*(*e*_1_).) Equation 3 says that predicting a higher-quality environment (i.e., the expectation of *e*_1_ or *E*(*e*_1_)) reduces the degree to which an organism adopts a phenotype suited for a low-quality environment.

The second proposition captures steps 3 to 4:

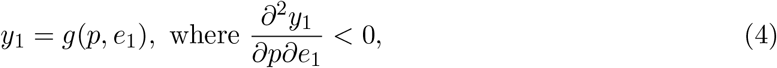

where *y*_1_ is the trait of interest, measured in adulthood. Note that *y*_1_ must be a trait that can be ordered to represent better or worse outcomes (e.g., good versus poor health). Equation 4 posits that adopting a phenotype in anticipation of a low-quality adult environment yields worse outcomes as adult environmental quality improves, when all else is held constant. Together, these two propositions (Eq. 3, 4) formalize steps 2 to 4.

An example can clarify the logic of the last two paragraphs. Suppose that low-quality environment (low *e*_0_) maps onto a cold environment, and that growing a thick coat (*p*) is a phenotypic adaptation to the cold environment. An organism that predicts that the adult environment will be warm (higher *E*(*e*_1_), as predicted by high *e*_0_), will grow a thinner coat (less fur *p*). If the adult environment is indeed warm as predicted, then this adaptation (less fur) is likely to improve adult outcomes (e.g., less fur *p* means less overheating, so higher *y*_1_).

Please refer to the supplement for related information about other variants of PAR (e.g., internal PAR, Section A.1), as well as information about the relationship between PAR and the developmental adaptive response hypothesis. This hypothesis proposes that an organism adapts to its developmental, rather than its predicted adult, environment (see Section A.2).

#### 2.2.1 Tests for PAR

Now that we have defined PAR, we can explicate the tests that are implied by the definition. There are two basic strategies: phenotypic adaptation tests and mismatch tests. Both of these strategies are appropriate for our mathematical models, and are regularly used in the literature [e.g. 32, 33].

The first strategy—phenotypic adaptation tests—focuses on the *mechanism* of PAR, i.e., the phenotypic “choice” an organism makes during development. We refer to the phenotypic adaptation (e.g., adopting a faster life history speed or developing a specific wing spot pattern [32, 34]) as a mechanism because developmental “choices” about these phenotypes are how an organism gets from predictions to an optimal fitness outcome. To understand the specific tests implied by this phenotype-focused strategy, we can collapse the three equations that capture PAR (Eqs. 2, 3, and 4) above into two expressions. The first is obtained by plugging steps 1 to 2 of our causal chain (Eq. 2) into steps 2 to 3 (Eq. 3) to obtain

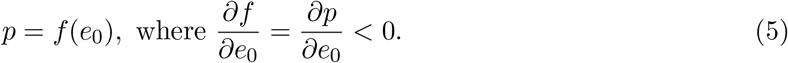

This inequality says that low-quality developmental environments cause phenotypic adaptations to low-quality adult environments. The inequality reflects steps 3 to 4 in our PAR causal chain (i.e., Eq. 4), which says that a phenotypic adaptation to a low-quality environment improves adult outcomes in that environment. These two expressions (Eqs. 4 and 5) imply two necessary tests that make use of data on organisms’ phenotypic adaptations. The first test, based on equation 5, asks whether the expected phenotype develops in response to conditions during development (making phenotype *p* the outcome variable and environment the regressor variable). The second test, based on equation 4, asks whether this phenotypic adaptation causes better health or fitness outcomes (making phenotype *p* now the regressor).

The second strategy for testing for PAR—mismatch tests—ignores information about specific phenotypes. Mismatch tests specifically evaluate whether an organism does better, on some health or fitness measure, if its adult environment is more similar to its early environment than it does if its early and adult environments are less similar (i.e., the mismatch hypothesis) [e.g., 20, 33]. This strategy plugs the inequality from Eq. 5 into steps 3 to 4 (Eq. 4) to connect developmental and adult environment, without reference to any specific phenotypic adaptation:

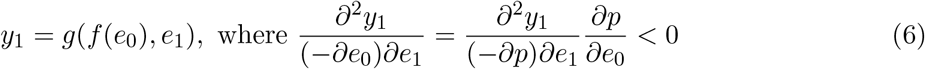

This inequality gives both the definition and the test of the mismatch hypothesis. To see why, it helps to evaluate the function at the point where *e*_0_ = *e*_1_. A concomitant decrease in developmental environmental quality and an increase in adult environmental quality (or vice versa) increases the mismatch between developmental and adult environments, leading to worse outcomes.^2^ This definition of the mismatch hypothesis is awkward because it relies on a cross-derivative and perhaps restrictions on *e*_0_, *e*_1_ (see note 2). Therefore, we simplify the mismatch hypothesis as follows:

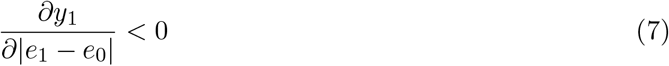

Now, the hypothesis states that adult outcomes decline as the gap between developmental and adult environments increases.

The first strategy—tests of phenotypic adaptation—can complement mismatch tests by providing more specific information about the mechanism(s) by which organisms arrive at outcomes. For more on this strategy, refer to the supplement (Section A.3). In the main text we will focus on mismatch tests because they are commonly invoked in the literature [e.g., 26, 20, 33].

### 2.3 Adult environmental quality (AEQ) hypothesis

Finally, we define the AEQ hypothesis, which says that a higher-quality adult environment will result in better adult outcomes, i.e.,

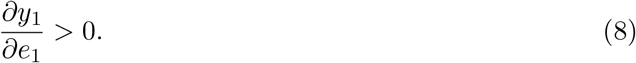

The AEQ is not a primary focus of the literature on understanding the early life drivers of inequality, as DC and are PAR are. We include it here to facilitate the discussion of conceptual issues surrounding all three of these hypotheses in the next section.

## 3 Conceptual issues in testing models

Four conceptual issues are central to any empirical tests of the DC, PAR, and AEQ hypotheses. We discuss two of these issues below. The other two (overlapping predictions generated by the theories, and the difficulty of testing intra-individual theories with inter-individual data) are discussed in Section A.4 in the supplement.

### 3.1 Issue 1: Non-independence of e_0_, e_1_, and ∆e

Because ∆*e* depends on *e*_0_ and *e*_1_, independently testing the DC, PAR, and AEQ hypotheses is challenging. Testing DC depends on variation in *e*_0_, testing PAR depends on ∆*e*, and testing AEQ depends on *e*_1_. Holding one of these quantities constant will unavoidably affect the others. For instance, if we hold *e*_0_ constant but increase *e*_1_, one cannot determine if observed changes in outcomes are associated with an increase in *e*_1_ or an increase in the change in environment ∆*e*. This means that researchers testing for DC and for PARs must make additional assumptions to also test the AEQ hypothesis. For example, one must define PAR as having equally deleterious effects regardless of whether the adult environment is better or worse than the developmental environment (“symmetric” rather than “asymmetric” PAR; see Section A.5 and Figure 3 in the supplement). Because symmetric PAR posits that changes in environment have the same effect on outcomes whether the change is positive or negative, a smaller reduction in outcomes with positive changes than with negative changes suggests AEQ, but also could, in theory, be consistent with an asymmetric PAR.^3^

To deal with this issue (an identification problem in statistics and economics literature parlance) in a way that does not arbitrarily prioritize testing one hypothesis over another, we recommend (i) explicitly stating the relationships among developmental environment, adult environment, and the difference between the two, and (ii) deciding which of these is/are independent of the theoretical model (i.e., exogenous to the model’s structure), and which are a byproduct of the model’s structure (i.e., *not* independent of the model’s structure, but simply a byproduct of the exogenous variation). Only hypotheses that concern the independent variation are testable [36].

The process that we feel best represents the phenomenon of interest here is that *e*_0_ and ∆*e* are independent, and that these collectively generate *e*_1_ via the formula *e*_1_ = *e*_0_ + ∆*e*. An alternative assumption—that ∆*e* is simply a byproduct—seems less defensible, because time moves linearly: *e*_0_ and ∆*e* come prior to *e*_1_, so it is unlikely *e*_1_ causes either *e*_0_ or ∆*e*. Imagine an adult baboon living through a drought. She has already experienced *e*_0_, and is now experiencing something (drought) that is familiar (i.e., matches her prior experience, a low ∆*e*), or unfamiliar (i.e., does not match her prior experience, a high ∆*e*). She does not experience *e*_1_ independent of what she experienced during development. One could assume a different process, but it is still the case that only two of the three hypotheses will be testable.

#### 3.1.1 Issue 2: Non-mutually exclusive theories

A second difficulty is that the DC, PAR and AEQ theories, as we have chosen to define them, are not mutually exclusive. Any combination of them could be true at the same time. This means that univariate models without higher-order polynomial terms (e.g., interacting and squared variables) may not be able to distinguish which theories are true. Consider the following simple model:

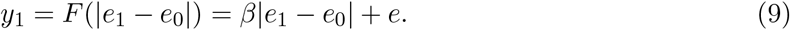

This model seeks to correlate adult outcomes with the mismatch between early life and adult environments. We differentiate this model from the theoretical models in the previous section by using capital letters to define the function. Suppose, however, that both DC and PAR are true. The model described as above (Eq. 9) does not simply control for or eliminate the influence of DC just by excluding a separate *e*_0_ term. Therefore, the estimated coefficient will suffer omitted variable bias and not equal the partial derivative of adult outcomes with respect to mismatch (*∂y*_1_*/∂*|*e*_1_ *− e*_0_|):

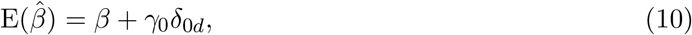

where *γ*_0_ is equal to *∂y*_1_*/∂e*_0_ and *δ*_0*d*_ is the coefficient from a regression of |*e*_1_ *− e*_0_| on *e*_0_. A similar problem afflicts any empirical specification that does not allow all plausible models to be true. At best, effect sizes are likely to be over or under-estimated; at worst, such models will generate answers that are the opposite of the real-world truth (e.g., concluding that PAR or DC do not exist when in fact they do).

## 4 Issues with a common visualization and testing strategy

One common empirical approach for implementing a mismatch test for external PARs (Section 2.2) relies on estimating a regression with interaction effects between developmental and adult environmental quality [e.g. 20, 33, 37, 38, 39, 40, 41], i.e.,

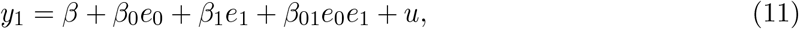

where *u* is a regression error term. We refer to this equation as “the interaction regression.” See Section A.3 in the supplement for a discussion of the interaction regression as it pertains to internal PARs [15].

Testing for DC with the interaction regression focuses on the coefficient on *e*_0_, while the test for a PAR focuses on the coefficient on the interaction term *β*_01_. This strategy is partly motivated by visualizations like the one depicted in Figure 2, which plot adult outcomes against adult environmental quality separately for an organism that experienced different developmental environments [e.g., 14, 20]. Failing to reject the hypothesis that *β*_01_ = 0 is interpreted as evidence for DC, while rejecting the hypothesis that *β*_01_ = 0 is interpreted as preliminary evidence for a PAR (so long as *β*_0_ *<* 0 is also true, as in Figure 2a; regardless of what *β*_0_ is, as in Figure 2b).

**Figure 2:**
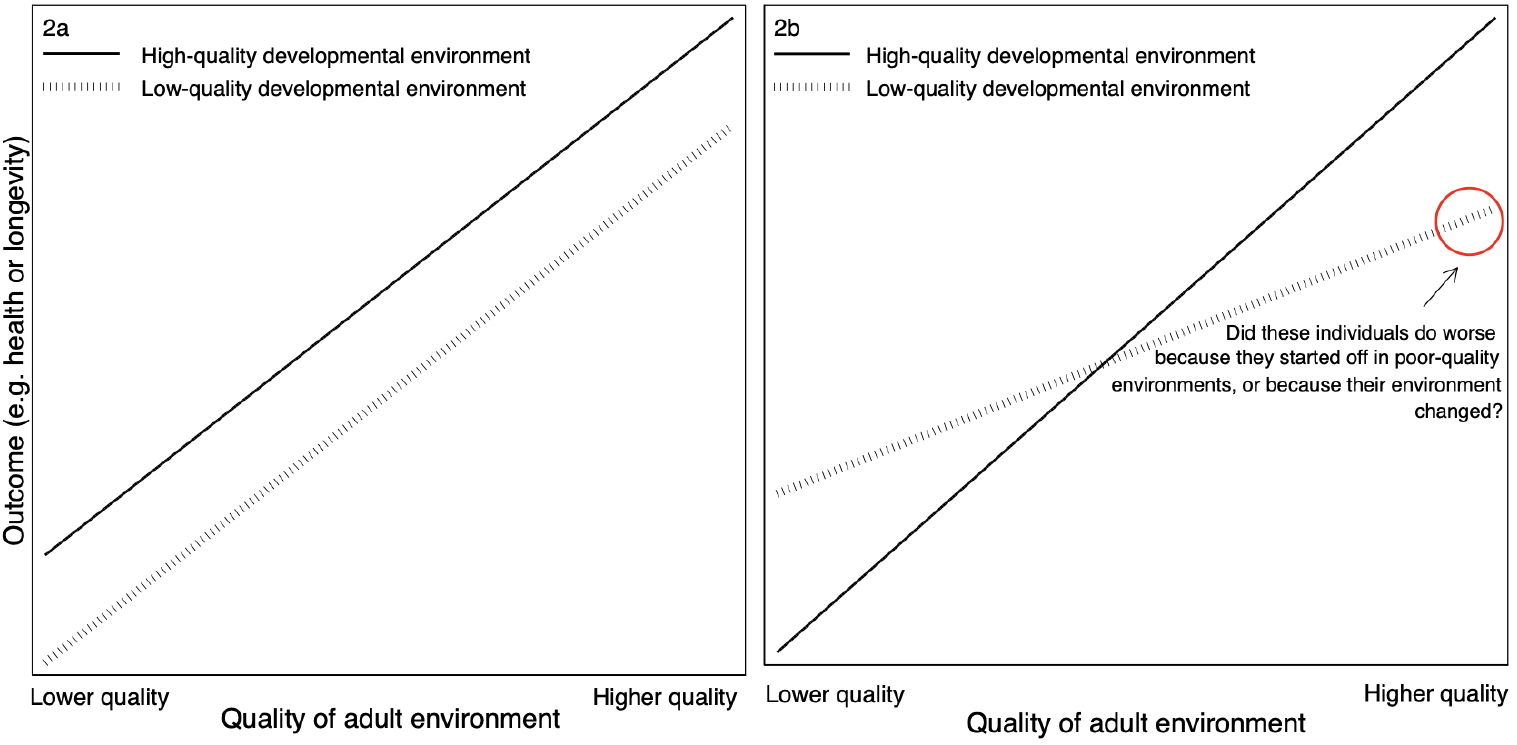
Commonly-used depictions of empirical evidence for the developmental constraints (DC, Section 2.1) and predictive adaptive response (PAR, Section 2.2) hypotheses. In panel **2a**, as adult environmental quality improves, health and fitness outcomes improve. However, organisms who started in low-quality developmental environments always fare worse than peers who started in high-quality environments. In panel **2b**, organisms that experienced “matching” (i.e., similar-quality) developmental and adult environments have better outcomes than organisms that experienced “mismatched” developmental and adult environments. These depictions manipulate both the starting point (*e*_0_ in Table 1, the variable developmental constraints theory is concerned with) and how well the developmental and adult environments match (∆*e* in Table 1, the variable predictive adaptive response theory is concerned with) simultaneously, making it difficult to distinguish between these two hypotheses. Moreover, the x-axis is the variable that the adult environmental quality hypothesis tests, i.e., that adult environmental quality (*e*_1_ in Table 1) impacts adult outcomes, holding all else constant.

To map the sign on the interaction term to a positive or negative effect of a change in environment, researchers often use visualizations to interpret the direction of *β*_01_. In these visualizations, if the line for organisms with high-quality developmental environments crosses from below the line for organisms with low-quality developmental environments (as in Figure 2b), this is interpreted as evidence for a PAR, because organisms whose developmental and adult environments “match” do better than those whose environments do not match.^4^ Figure 2b is indeed consistent with predictions derived from the PAR hypothesis, because organisms do better in adult environments that are more similar to their developmental environments. It is also consistent with the AEQ hypothesis (8), since all organisms do better as their adult environmental quality improves.

However, any visualization with the axes in Figure 2 is manipulating two variables simultaneously—the developmental environment (implicitly, DC) and changes between the developmental and adult environment (PARs). This makes it hard to determine which one is responsible for the observed effect. For example, on the right-hand sides of Figures 2a and 2b (where adult environmental quality is high), the organisms represented by the dashed line began their lives in a low-quality environment, which may account for their worse outcomes relative to the organisms who began their lives in a high-quality environment (solid line). However, the organisms who began in a low-quality environment are *also* in an adult environment that does not match their developmental environment (i.e., they experienced change), while the organisms who began in a high-quality adult environment are in an adult environment that matches their developmental environment (i.e., they did *not* experience change). It is not clear whether the individuals with high-quality developmental environments have better outcomes because they started their lives on top, or because their developmental and adult environments match.

Comparing individuals from low and high-quality developmental environments in low-quality adult environments (the left-hand side of the x-axis in Figures 2a and 2b) still does not cleanly separate the effects of starting point from the effects of mismatch, because it is still manipulating two different variables while embedding information about a third (adult environmental quality). Interpreting the “crossover” as a PAR when adult environmental quality is poor requires the assumption that the AEQ is true. It may well be, but it is far better to make this assumption explicit, and even better to test it (presuming that one has decided the relationship is testable; see Section 3.1). Fundamentally, the visualization is flattening a three (or higher) dimensional problem into two dimensions, and thus conflating the two primary variables of interest. See the supplement for more details (A.6).

Using adult environmental quality—the variable of interest in the AEQ hypothesis—as the x-axis variable generates interpretability problems when testing DC or PAR. When visualizing DC, we recommend creating figures with an x-axis that represents the developmental environment (Figure 1a), and when visualizing PARs, creating figures with an x-axis that reflects how well the developmental and adult environments match (Figure 1b and 1c). This allows allows for separate visual identification of DC and/or PARs, as well as determination of how a negative change in environment might be different than a positive one (see also [33]).

## 5 Constructing theoretically-derived tests of the DC and PAR models

Using an interaction regression to test for PARs is problematic [and is something we ourselves have done: 32, 33]. In order to demonstrate why, we will first describe the empirical tests that follow from our formal definitions of DC and PARs.

The definitions of DC and PARs are compatible with a general functional form:

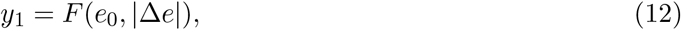

where capital *F* indicates an empirical function involving observed data. We do not include adult environment in the function (and thus the AEQ hypothesis) for the reasons given in Section 3.1.

Testing these theories with data requires a functional form or specification that we can estimate, for example,

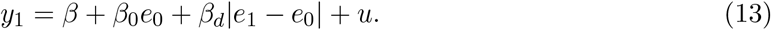

where *β*_*d*_ is the marginal effect of the absolute value of the mismatch between developmental and adult environments.

### 5.1 Regression model derived from mathematical definitions

To obtain a functional form, a reasonable approach is to assume *F* is differentiable over the relevant range and take a Taylor expansion, which yields a power series. We choose a second-order expansion around *e*_0_ = *e*_1_ = |∆*e*| = 0 to limit the number of terms that must be estimated. Estimating a regression model with only first-order terms cannot capture both DC and PAR. Moreover, it would be difficult to relate to the interaction regression (Eq. 11), since an interaction term requires a second order (or higher) expansion. A higher than second-order expansion is possible, but requires more data to be adequately powered relative to a model with fewer terms. Sample sizes in the real world often cannot accommodate this requirement. A second-order expansion implies the following regression model:

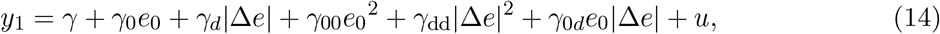

where the subscript 0 on *γ* indicates the coefficient is on *x*_0_, *d* indicates it is on |∆*e*|, 0*d* indicates it is on *e*_0_|∆*e*|, and so on. The error *u* captures omitted, orthogonal factors that influence the observed outcomes in a given dataset. The first three terms are included in a first-order Taylor expansion; the last three terms, including the interaction term (Eq. 11), are added by a second-order expansion.

Equation 14 is the primary regression model that we recommend and evaluate in the remainder of this paper. It balances capturing elements omitted from the interaction model (which we discuss in the next section) with power limitations of real-world data sets. Researchers with larger data sets may attempt to estimate higher order polynomials and apply the appropriate tests implied by the definitions of DC and PARs.

### 5.2 Testing for DC and PARs with a quadratic regression

The quadratic regression (Eq. 14) implies specific and different statistical tests for DC and PARs. The definition of DC (Eq. 1) implies the following test for DC:

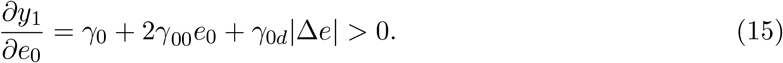

If one cannot reject that *∂y*_1_*/∂e*_0_ = 0 in favor of the inequality above, then one cannot reject the null hyopthesis that DC is false. Moreover, the prediction of the PAR hypothesis in equation 7 implies the following test for PARs:

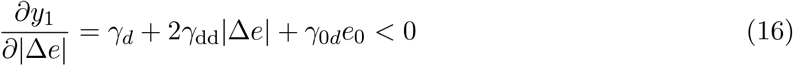

If one cannot reject *∂y*_1_*/∂*|∆*e*| = 0 in favor of the inequality above, then one cannot reject the null that PAR is false.

## 6 Implications for the interaction regression model

Although the interaction regression (Eq. 11) is commonly used to test for PARs, and also may be used to test for DC (via the coefficient on *e*_0_), we have identified several problems with using that regression to test these hypotheses. In this section, we show why the interaction regression is mathematically incompatible with the existence of PAR as we formally define it. In Section A.7 we also show why the test for DC in an interaction model may yield a different result than the simple test for a correlation between developmental environment and adult outcomes, and why, if the interaction regression does not accurately describe reality, the coefficients estimated from that regression are likely to be misleading.

If the interaction regression were a correct description of reality, the mismatch prediction of PARs could not be true. To see why, we plug *e*_1_ = *e*_0_ + ∆*e* into the interaction model (Eq. 11) so we can take the derivative of the latter with respect to ∆*e*. Using the chain rule, we can now take the total derivative of the interaction model with respect to |∆*e*|:

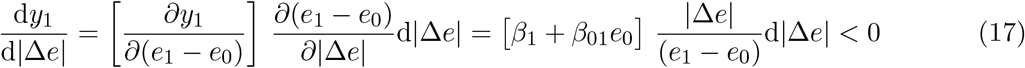

for positive d|∆*e*|. The prediction of the PAR hypothesis in (7) implies that PARs exist if this derivative is negative. Note that the derivative must be negative when ∆*e* is both positive <u>and</u> negative. This is true even if the environment is coded as a binary variable. For example, coding low as *e* = 0 and high as *e* = 1 yields mismatch if ∆*e* is 1 or *−*1 or, equivalently, |∆*e*| *>* 0. Regardless of how environment is coded, if ∆*e >* 0, then ∆*e/*|∆*e*| *>* 0, and if ∆*e <* 0, then ∆*e/*|∆*e*| *<* 0. So, for the inequality in equation 17 to hold, *β*_1_ + *β*_01_*e*_0_ has to be negative for positive ∆*e* and positive for negative ∆*e*. However, if we start from a given developmental environment *e*_0_, this is impossible: *β*_1_ + *β*_01_*e*_0_ is constant and does not flip signs. Importantly, the mismatch prediction of PARs are not a theoretical impossibility with the quadratic regression because it includes a |∆*e*| term; the derivative in the quadratic regression (Eq. 16) can theoretically be negative for the full range of ∆*e*.

Strictly speaking, the interaction regression will only fail to find evidence of PARs if researchers test for PARs based on the prediction in equation 7 (adult outcomes decline as the gap between developmental and adult environments increase). The interaction regression cannot generate results that pass that test. However, this impossibility likely also undermines the two-part test that supplements the interaction regression with a visualization of the interaction. That two part-test can be thought of as an effort to approximate testing the prediction in equation 7, but if directly testing that prediction with the interaction regression cannot find evidence for mismatch, then the approximation that combines the interaction regression with a visualization is also unlikely to find it.

## 7 Simulations of alternative tests for DC and PARs

We conducted simulations to determine how often the interaction regression (Eq. 11), as compared to the quadratic regression (Eq. 14), correctly finds or rejects the predictions generated by DC or PAR. Our process has five steps. We sketch our design below, but provide additional details in the supplement (Section A.9).

### 7.1 Simulation design

#### Step 1: Generate a large number of different virtual realities

Each simulation posits a virtual reality where, by assumption, a 3rd-order polynomial perfectly describes the effect of: (a) an organism’s developmental environment and (b) the mismatch between their developmental and adult environments on adult outcomes. A 3rd-order polynomial is the lowest-order polynomial that allows both the interaction and quadratic regression models to generate erroneous test results and compare error rates.

A 3rd-order polynomial can have infinitely different coefficient values and thus describe infinitely different realities. We pared those possible realities down by only allowing 5 evenly spaced values of *e*_0_ *∈* [0, 1] and of ∆*e ∈* [*−*1, 1], and by ruling out realities where at any allowed value of *e*_0_ or ∆ generated an outcome outside the range of [0, 1]. This left 130,201 “pruned” realities where each reality is defined by a 3rd order polynomial and a 10 × 1 vector of parameters for that polynomial.

#### Step 2: Determine whether PAR or DC are true in each reality

Each of the feasible realities was evaluated for PARs and DC by applying the tests in equations 7 and 1, respectively. Of all the pruned realities, 2,697 (2.07%) were truly positive for PARs, 2,697 (2.07%) were truly positive for DC, and 58 (0.04%) were truly positive for both. These feasible and true positive realities are the benchmark against which tests based on each regression model are evaluated.

#### Step 3: Simulate a data set for each reality

For each pruned reality, we generated a simulated data set for estimating regressions by adding noise to the outcomes that are generated in that reality for 16 different combinations of developmental and adult environments. Each data set included 2000 observations on the variables (*ŷ*_1_, *e*_0_, ∆*e*). For each observation, *e*_0_ is drawn uniformly from 4 evenly spaced points between 0 and 1; ∆*e* from a normal distribution with mean *−*0.03 and standard deviation 0.21^5^ but truncated at −1 and 1; and an error term *v* from a normal distribution with mean 0 and variance equal to that of the outcome at the mean value of (*e*_0_, ∆*e*) in each reality. The observed outcomes *ŷ*_1_ is generated by adding *v* to the true outcome in that pruned reality at the drawn (*e*0, ∆*e*).

#### Step 4: Apply different empirical tests for PAR and DC on simulated data for each reality

We apply tests that researchers currently use (based on on the interaction regression) and tests we recommend (based on the quadratic regression) to determine if each test finds that there is PAR and DC in each virtual reality. Specifically, we generated four test results for PARs in each reality:

1. **PAR test 1: Visualization test for PARs using the interaction regression (Eq. 11)**. This test finds evidence for PARs if (i) *β*_01_ ≠ 0, and (ii) the visualization shows that the fit line depicting the adult environment/adult outcomes relationship for organisms from low-quality developmental environments (the dotted line in Figure 2b) *intersects from above* the same line for organisms from high-quality developmental environments (the solid line in Figure 2b).
2. **PAR test 2: “Relaxed” version of the visualization test with the interaction regression (Eq. 11)**. In the presence of development constraints (DC), the fit line for adult environment/adult outcomes for organisms from low-quality developmental environments may be shifted downwards relative to organisms from medium- and high-quality environments (as depicted in Figure 1c). In this case, PARs might exist even if the lines for low- and highquality developmental environments do not cross. This suggests a relaxed visualization test which finds evidence for PARs if 1) *β*_01_ ≠ 0, and 2) the visualization shows that the fit line depicting the adult environment/adult outcomes relationship for organisms from lowquality developmental environments has *a lower slope* than the same line for organisms from high-quality developmental environments.
3. **PAR test 3: Theoretically-motivated test (Eq. 7) applied to the interaction regression (Eq. 11)**. This test finds evidence for PARs if the derivative of the interaction regression with respect to |∆*e*| is negative (i.e., Eq. 17). We implement this test notwithstanding the fact that the PAR hypothesis cannot be true if the interaction regression is a correct specification of reality.
4. **PAR test 4: Theoretically-motivated test (Eq. 7 applied to the quadratic regression (Eq. 14)**. This test finds evidence for PARs if the derivative of the quadratic regression with respect to |∆*e*| is negative (Eq. 16).

We also generated three tests for DC in each reality:

1. **DC test 1: Naive test for DC with the interaction regression (Eq. 11)**. This test finds evidence for DC if *β*_0_ in equation 11 is positive.
2. **DC test 2: Theoretically-motivated test (Eq. 1) applied to the interaction regression (Eq. 11)**. This test finds evidence for DC if the derivative of the interaction regression with respect to *e*_0_ is positive, i.e., *β*_0_ + *β*_01_*e*_1_ is positive.
3. **DC test 3: Theoretically-motivated test (Eq. 1) applied to the quadratic regression (Eq. 14)**. This test finds evidence for DC if the derivative of the quadratic regression with respect to *e*_0_ is positive. This test is presented in equation 15.

### 7.2 Evaluating empirical tests

The final step is to compute the rate at which each test correctly concludes there is PAR or DC and correctly concludes there is not PAR or DC. The top panel of Table 2 provides the sensitivity and specificity of each of the four methods of testing for PARs across all feasible realities (sensitivity is the percent of the 2,755 realities where PAR was true in which PAR was correctly detected; specificity is the percent of the remaining 127,504 realities where PAR was not true in which it was correctly <u>not</u> detected). The second column shows the performance of a coin-flip test, which we use as a benchmark of a data-uninformed test (i.e., a test with performance equal to chance).

**Table 2:**
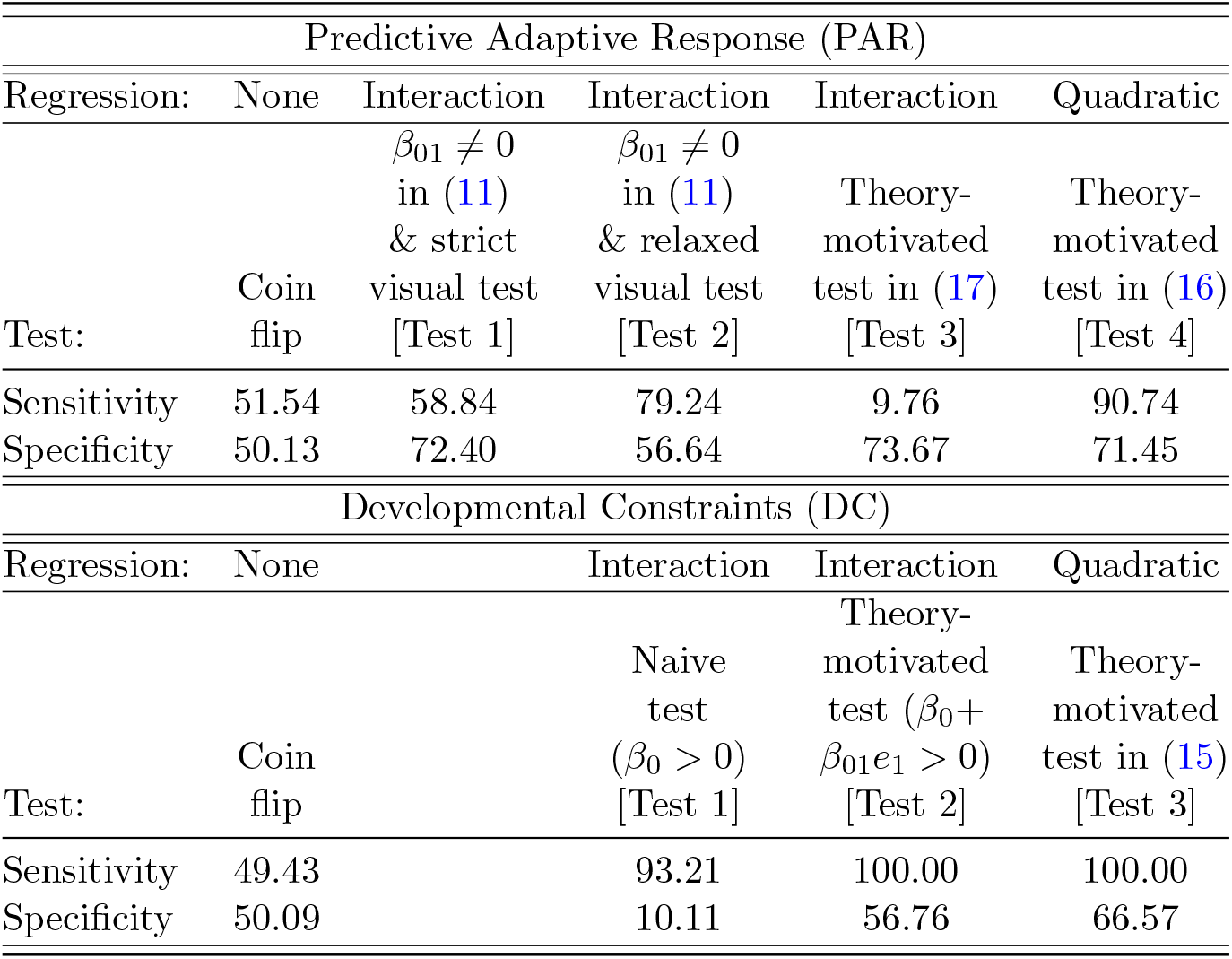
Percent of simulated realities where PARs and DC were correctly detected (sensitivity) and correctly not detected (specificity) using different tests.

The interaction regression has poor or imbalanced performance across a range of tests. For instance, the interaction regression with the visualization approach to testing for PARs has sensitivity that is only slightly better than a coin flip (58.84%). A relaxed visual test has higher sensitivity (79.24%) but lowered specificity, only marginally better than coin flip (56.64%); in other words, it incorrectly detected PAR 43.46% of the time. The interaction regression showed even worse performance with the theoretically-motivated test for PAR: sensitivity (9.76%) was much worse than a coin flip. This very poor sensitivity is to be expected: if the interaction regression is a correct description of reality, we demonstrated that one theoretically cannot find PAR. The visualization test using the interaction regression actually gets the right answer more often than the theoretically-motivated version precisely *because* the former is not actually testing the prediction generated by the PAR hypothesis (Eq. 7). This allows it to perform the same as or marginally better than a coin flip, while a theoretically-motivated use of the interaction regression is specifically biased *against* finding PARs even when they exist.

The quadratic regression combined with a theoretically-motivated test performs best of all. Sensitivity (90.34%) is higher than any test using an interaction regression and specificity is roughly the same (71.61%) as the best tests under the interaction model. This test is not perfectly sensitive and specific because it too suffers from bias due to omitted variables: that is, the realities it approximates also have third-order terms.

The bottom panel of Table 2 provides the sensitivity and specificity of three methods of testing for DC across feasible realities. The interaction model performed poorly relative to the quadratic regression. The naive test using the interaction regression has good sensitivity, but much worse specificity than a coin-flip. The test often finds DC whether it is true or not. The interaction regression combined with a theoretically-motivated test has perfect sensitivity, but specificity that is only marginally better than a coin flip. Switching to a quadratic regression and using a theoretically-motivated test for DC performs best of all. It too has perfect sensitivity, and somewhat better specificity (66.57%).

## 8 Discussion

DC and PAR are currently the most commonly invoked evolutionary explanations for early life determinants of inequality in adult health and fitness outcomes. However, these theories lack precise and consistent definitions. Further, different forms of DC and PAR make their own assumptions, assumptions that need to be made explicit rather than left implicit. Making assumptions and definitions explicit has the benefit of clarifying where different flavors of hypotheses do and do not generate differentiating predictions, what steps researchers must take in order to differentiate between them, and under what conditions it is possible for them to do so.

For example, the common notion that phenotypic adaptation tests (Section 2.2.1) are tests of an internal PAR and mismatch tests are tests of external PAR are not defensible, if the only distinction between the theories is the type of cue that organisms are using (which is simply a re-definition of the *e*_0_ and *e*_1_ terms in Table 1); this is made apparent by the use of formal definitions and explication of definitionally-aligned tests. Another example—and one with potentially major ramifications—is the distinction between intra-versus inter-individual *theories* as opposed to intra-versus inter-individual *tests* (Section 2). DC and PAR are theories about the causal effects of environments on individuals. Methodological limitations mean that it is rarely possible to observe the same individual in different environmental conditions. Being forced to rely on inter-individual tests requires us to make strong assumptions about the distribution of individuals across environments in order to draw meaningful conclusions about an intra-individual theory. This statistical problem is beyond the scope of this paper, but it is a critical example of how removing definitional ambiguity and explicating assumptions can improve the science of early life effects.

However, in this paper we have chosen to focus on one specific consequence of the lack of formal definitions for DC and PAR, namely, that a common test meant to disentangle these hypotheses— linear regression models with interaction effects—ends up conflating the two theories. Testing one of the key predictions generated by the PAR hypothesis requires researchers to detect the effects of environmental mismatches. Doing so with any accuracy is difficult with an interaction model and its complementary data visualization strategy. Indeed, such an interaction model is theoretically incapable of testing the mismatch prediction of PAR. A test that applies the mismatch prediction to the quadratic regression does not suffer from a basic mathematical incompatibility problem. Similar arguments justify the use of a reasonable, formal definition of DC along with a quadratic regression. Our simulations show that using the theoretical predictions of PARs and DC specified in Table 1, together with the quadratic regression, dramatically improved sensitivity/specificity trade-offs (relative to any use of an interaction model) when testing for PARs and DC. Critically, there is no downside to using the quadratic specification, beyond requiring slightly larger sample sizes. It will produce reliable answers even in a world where an interaction regression is an accurate representation of the biological reality, a possibility we discuss in more detail below. The reverse is not true, for the many reasons discussed above.

Currently, support for PARs in the literature is mixed, especially in mammals [11]. The definitional and testing issues highlighted here raise the question of whether this is because of flaws in the methods used to detect them. Our results show that statistical tests derived from mathematical definitions of the DC and PAR concepts, along with more flexible regression models, provide clearer answers, and will improve our ability to compare results across studies.

Because there is already a literature that employs visualizations such as Figure 2 and the interaction regression in equation 11 to test for PARs, it is useful to know if there are conditions under which this approach is a valid test of the PAR hypothesis. Sufficient conditions for the interaction regression to be valid are if: (a) mismatches in only one direction reduce health/longevity/fitness, *and* (b) the relationship between outcome, developmental environment and the change in environment over a lifetime is quadratic, but with certain parameter restrictions that cause the quadratic model to be identical to the interaction model. For example, focusing on a positive change in environment, the second condition is that the following are true: (i)

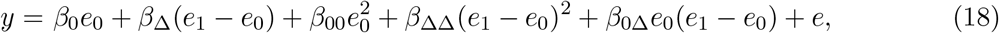

and (ii) *β*_∆∆_ = 0, *β*_00_ = *−β*_0∆_, and *β*_0∆_ ≠ 0. Under these parametric restrictions, the quadratic regression above collapses to something like the interaction regression in equation 11. Of course, it is difficult to know *ex ante* whether the second condition is satisfied without first estimating a quadratic regression. Given the high rates of both false positives and false negatives generated by the interaction model plus visual test strategy (Table 2), it is reasonable to infer that somewhere between 20 and 40% of the PAR-related results in the literature are false negatives, 30-45% are false positives, and that it may contain a significant number of false positives for DC. Of course, this assumes that researchers are equally likely to publish null results and results in which they found evidence for PAR and/or DC. Assuming the “file drawer” effect applies to this literature [43], then false positives may be over-represented relative to false negatives.

The DC and PAR hypotheses appear in the literature of many academic fields [4, 15, 16, 18, 33, 35]. Significant human and financial resources are being devoted to untangling their effects because of their important implications for public health and policy, and for our understanding of if and how variation in early environments explains inequality in adult outcomes. Clear, careful definitions and appropriate statistical tests are absolutely essential for forward progress in this important research area. We recommend that researchers take the following steps when testing the DC and PAR hypotheses:

1. Rely on statistical tests that are derived from mathematical definitions, to avoid conflating different phenomena being evaluated in the same model.
2. Avoid using interaction or first-order polynomial models to test for environmental mismatch effects; instead, use a quadratic or higher-order regression model. We provide Stata and R code at (github link) to assist with implementation of quadratic models and the associated statistical tests for DC and PARs.
3. Verify that data visualizations cleanly separate the concept(s) of interest. It is better to use multiple visualizations that each address a single phenomenon than a single visualization that potentially conflates different phenomena.
4. Keep in mind that the overlapping predictions made by DC and PAR hypotheses and the non-mutually exclusive nature of these hypotheses mean that separately identifying the effects of early, adult, and mismatched environments is usually not possible.

## 9 Acknowledgements

We thank A. Lea, C. Weibel, B. Lerch, N. Grebe, F. Campos, and especially S.C. Alberts and J. Tung, whose insights greatly improved the manuscript. We gratefully acknowledge the support of the NIH, especially the National Institute on Aging, particularly NIH Grants R01AG053330, R01AG053308, R01HD088558, and P01AG031719. We also thank the members of the Amboseli Baboon Research Project (ABRP), which inspired our evaluation of these models. The ABRP is grateful to the Kenya Wildlife Service, Institute of Primate Research, National Museums of Kenya, National Council for Science and Technology, the Kajiado County Council, the members of the Amboseli–Longido pastoralist communities in Kenya, and the Enduimet Wildlife Management Area, the long-term field team (R. S. Mututua, S. Sayialel, and J. K. Warutere), and V. Oudu and T. Wango for their assistance in Nairobi. Malani acknowledges the support of the Barbara J. and B. Mark Fried Fund at the University of Chicago Law School.

## A. Supplementary materials

### A.1 Variants of predictive adaptive response

A prominent variant of the PAR hypothesis, internal PAR, proposes that organisms make predictions based on their somatic state, rather than the external environment [44, 15, 45, 32]. One can capture this version of PAR with steps 1-4 of our PAR causal chain (in Section 2 in the main text) by replacing either *e*_1_ only or both *e*_0_ and *e*_1_ with an organism’s internal state in the relevant life stage(s) [15]. While the internal PAR hypothesis typically proposes that organisms use their external environment during development to predict their somatic state in adulthood [15], internal state during development could also be used to predict internal state in adulthood. Somatic state could theoretically be substituted for external environment either at *e*_1_ only, or at *e*_0_ and *e*_1_. The testable predictions remain as in equations 5 or 7 in the main text, so long as an organism uses its developmental environment (or its developmental somatic state) to predict its future somatic state. A major challenge to distinguishing internal from external PAR, however, is that when internal states are influenced by external environments—the premise upon which the internal PAR theory is based—observed differences in the former may simply reflect unobserved differences in the latter. In the literature, tests for internal PAR have focused on phenotypic adaptations, while tests for external PAR have focused on environmental mismatch [15, 26] (see Section 2.2.1 for an overview of the differences in these testing strategies). However, if the definitions of the concepts are the same except for re-defining one or both of the *e* terms, this distinction has no theoretical justification. External PAR can be tested using a phenotypic adaptation strategy (Eq. 5), and internal PAR could be tested using mismatch (Eq. 7).

There are also other possible variants of PAR that one could define. One variant would allow an organism to predict a future environment that differs from its developmental environment (contra steps 1 to 2 of the causal chain that defines PAR in Section 2.2 in the main text). For example, humans may predict that our future environment will be warmer; a lynx born during a season with abundant hares to eat might predict (perhaps not consciously) an eventual crash in the hare population. This is simply *E*(*e*_1_) ≠ *e*_0_. More specific variants (i.e., that adult environment is going to be twice as good as developmental environment, or twice as bad) could easily be defined as well. Another variant concerns whether overly optimistic and overly pessimistic predictions have differential effects [14]. We will define “symmetric mismatch” as a version of mismatch wherein positive and negative values of ∆*e* = *e*_1_ *− e*_0_ have the same (negative) effect on outcomes:

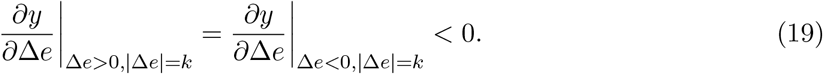

A visualization of the prediction this version would generate can be found in the left panel of Figure 3. By contrast, we will define “asymmetric mismatch” as a version of mismatch wherein positive and negative mismatch each have deleterious effects on outcomes, but equal-magnitude positive and negative errors do not have the same magnitude of deleterious effects on outcomes. A visualization of this prediction can be found in the right panel of Figure 3. These prediction errors could theoretically be asymmetric in the opposite direction; our choice to depict negative predictions as more deleterious than positive ones is arbitrary. The reason that these variants are important is because they affect whether the DC, PAR, and AEQ hypotheses can be empirically distinguished, a topic we discuss below; see Section A.5.

**Figure 3:**
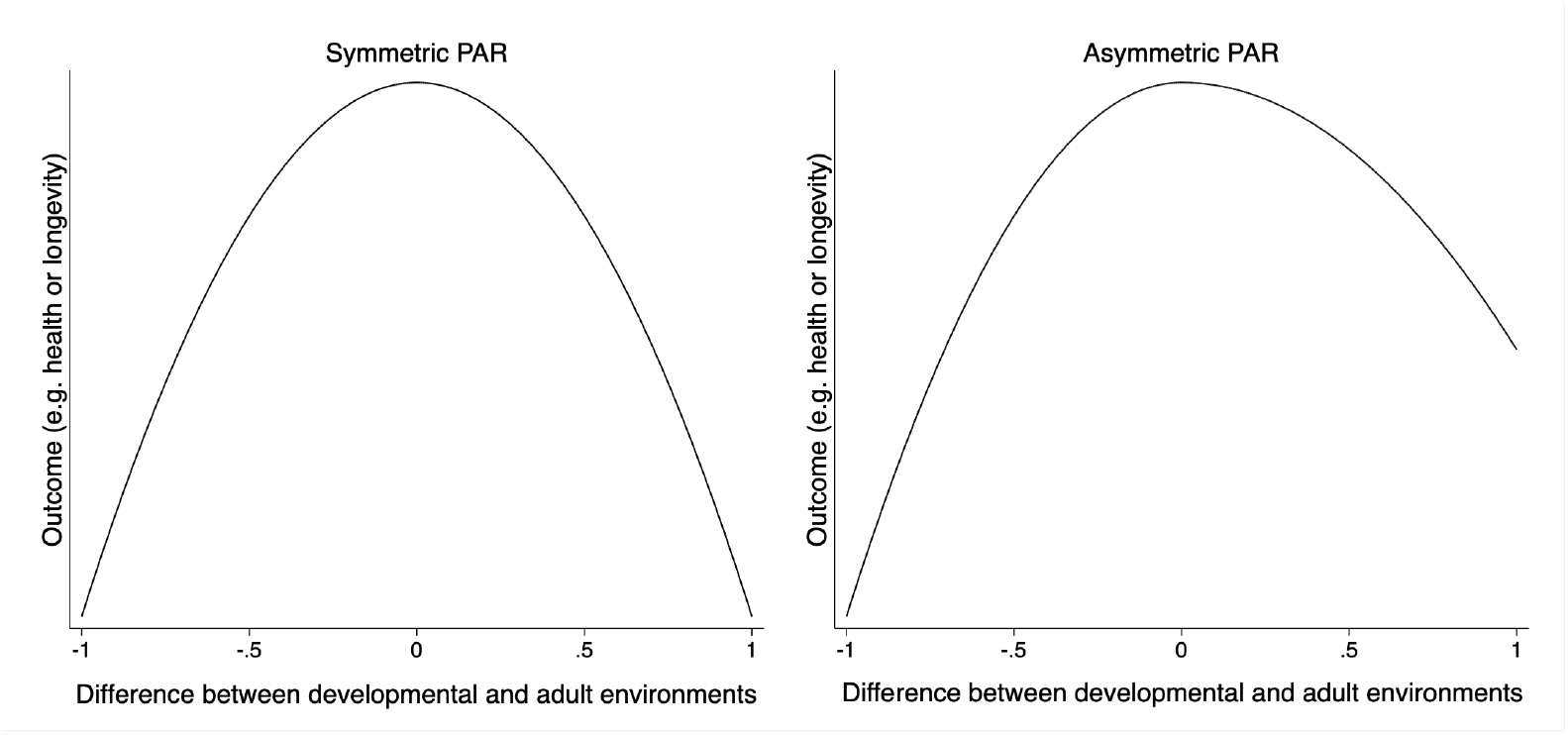
Visual depictions of the concept of symmetrical and asymmetrical predictive adaptive responses

### A.2 PAR and the developmental adaptive response hypothesis

The developmental adaptive response (DAR)^6^ model is theoretically distinct—but often empirically indistinguishable—from PAR. DAR posits that an organism adapts to its developmental environment not because that environment is its best prediction of its adult environment, but because its goal was simply to adapt to its developmental environment (Table 3). One example of a DAR comes from water fleas, who acquire a protective “helmet” when they receive signals that they will be in a predator-rich environment [47]. This is considered a DAR, rather than a PAR, because the fleas benefit immediately from their phenotypic adjustment [48]. This theory captures a common alternative explanation for observations that mismatched developmental and adult environments may lead to poor outcomes [17, 49, 11]. While DARs and PARs are theoretically distinguishable (and depending on circumstance, even mutually exclusive), DARs generate the same prediction as PARs, which makes distinguishing the theories difficult.

**Table 3:** Formal definitions and derived predictions of theories for the relationship between the quality of developmental environments and adult outcomes.

| Theory | Definition | Observable Variation | Prediction or Test (Eqn. No.) |
| --- | --- | --- | --- |
| Developmental Constraints (DC) | $\frac{\partial y_1}{\partial e_0} > 0$ | $(y_1, e_0)$ | $\frac{\partial y_1}{\partial e_0} > 0$ (1) |
| Predictive Adaptive Response (PAR) | $E(e_1) = e_0$<br>$\frac{\partial p}{\partial E(e_1)} < 0$<br>$\frac{\partial^2 y_1}{\partial p \partial e_1} < 0$ | | |
| Phenotypic adaptation test | | $(e_0, p)$ | $\frac{\partial p}{\partial e_0} < 0$ (5) |
| Environmental mismatch test | | $(y_1, \vert \Delta e\vert )$ | $\frac{\partial y_1}{\partial \vert \Delta e\vert } < 0$ (7) |
| Developmental Adaptive Response (DAR) | $\frac{\partial p}{\partial e_0} < 0$<br>$\frac{\partial^2 y_0}{\partial p \partial e_0} < 0$ | $(e_0, p)$ | $\frac{\partial p}{\partial e_0} < 0$ |
| Adult Environmental Quality (AEQ)* | $\frac{\partial y_1}{\partial e_1} > 0$ | $(y_1, e_1)$ | $\frac{\partial y_1}{\partial e_1} > 0$ (8) |

The causal chain that motivates DAR is described by the following three steps:

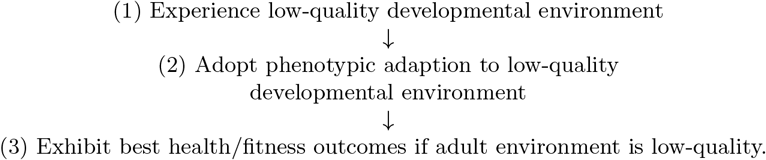

The only difference between DAR and PAR is the removal of the prediction step in the causal chain for PAR (i.e., step 2 in the PAR chain; see Section 2.2 in the main text). The organism is adapting to the developmental environment, which is directly observable, and thus no prediction is necessary.

Two mathematical formulas define DAR. Steps 1 and 2 of the DAR causal chain are captured by

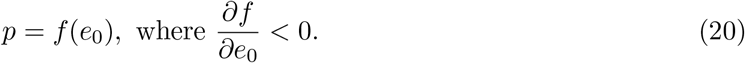

That is, phenotypic changes are a direct response to the developmental environment. The same is true for PAR, except that the developmental environment matters *indirectly* because it is used to predict the adult environment. Steps 2 and 3 for DAR imply

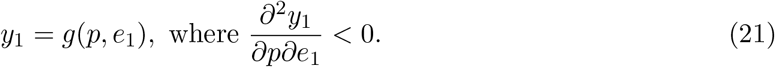

That is, phenotypic adaptations to a low-quality developmental environment harm adult outcomes when the adult environment is high-rather than low-quality. This claim is the same as steps 3 and 4 of the PAR causal chain.

The similarity between the equations for DAR (20, 21) and PAR (3, 4) highlight the difficulty of distinguishing DAR from PAR. DAR’s first proposition is identical to the first test for PAR in (5). If we plug the first DAR equation into the second, the result,

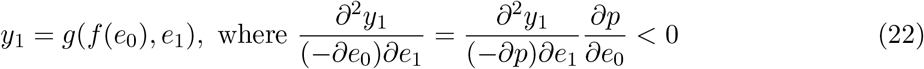

is identical to the mismatch hypothesis (6) that flows from PAR.

There are some circumstances in which it is possible to distinguish DAR and PAR. If the phenotypic adaptation for DAR is different than for PAR, i.e., one examines an adaptation that is only useful in early life (or only useful in later life), then one can test for DAR but not PAR (and vice versa). For example, locust wing phenotypes vary between adult morphs depending on the developmental environments larvae are exposed to, but the wing adaptations—which are not expressed until adulthood—clearly have no benefit during the larval phase [50, 51, 52, 53]. This approach to distinguishing DAR and PAR requires the use of a phenotypic adaptation test (discussed in Section 2.2.1 in the main text, and Section A.3 below). The mismatch test yields the same prediction for DAR and PAR because it does not examine phenotypic adaptations.

A primary goal of this paper is to address the conceptual framing of the literature on DC and PARs, so we refer to the prediction that a mis-fit between an adaptation and adult environment leads to worse adult outcomes as PAR rather than DAR, even though the two are empirically indistinguishable with a mismatch test.

### A.3 Testing for predictive adaptive responses using phenotypic adaptations and phenotype/environment interactions

As discussed in Section 2.2.1, there are two strategies for testing PAR. One is to look at the impact of differences between early life and adult environments, i.e., test the mismatch hypothesis. We discussed this strategy in depth in the main text. The other strategy is to examine the phenotypic adaptations to developmental environments (sometimes referred to as developmental inputs [26]) and the impact that these phenotypes have on the relevant outcome(s). We explore this second strategy here.

Recall that the main text noted that the 4 steps in the causal chain for PAR can be expressed as 3 equations, and that if one of the equations (for step 1) is plugged into another (for steps 2 to 3), one obtains a simplified expression for PAR with just two equations:

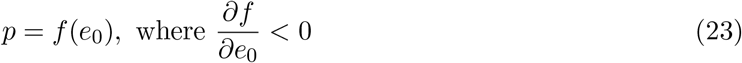

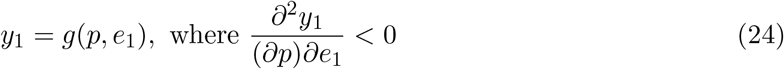

where *p* are phenotypic adaptions to low-quality environments. The first equation says that a low-quality developmental environment triggers phenotypic adaptations suitable for a low-quality adult environment (because the adult environment is expected to be the same as the developmental environment).

The second equation says that outcomes are better with the relevant adaptation if the adult environment is low-quality, than they would be if the adaptation had not occurred. While our statement of PAR talks about external environment, one could substitute internal somatic state for environment and the hypothesis would stand.

This simplified definition suggests two empirical tests for PAR that rely on observed phenotypic adaptations. One test uses the first equation (23) in the simplified definition, i.e., it checks if organisms from a low-quality developmental environment make a phenotypic adaptation to that environment. The most basic regression specification for this test is

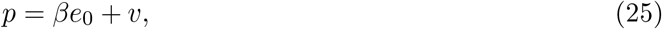

where the test for PAR is that *β <* 0. In the main text we recommend *against* specifications that employ first-order approximations for *y*_1_ = *f* (|∆*e*|) such as *y*_1_ = *γ*|∆*e*| + *e* due to the risk of omitted variable bias in estimates of *γ* (Section 3.1.1). That is not a problem for (25) because there are no other possible (known) triggers for an adaptation that is correlated with developmental environment. ^7^

A second empirical test for PAR that examines phenotypic adaptation focuses on the second equation (24) in the simplified expression of PAR. A second-order approximation yields a quadratic regression of the form

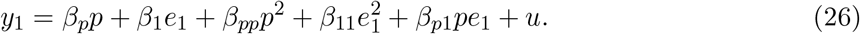

The test for PAR is that

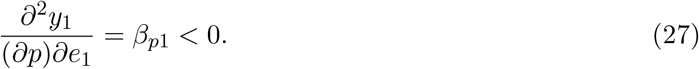

Given how simple this test is, one might be tempted to estimate an interaction model of the form:

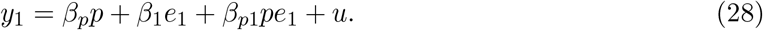

However, this would yield a biased estimate for *β*_*p*1_ because it omits squared terms *p*^2^ and 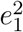, whichare correlated with the interaction term *pe*_1_.

[26] has suggested an empirical analysis that somewhat resembles the preceding interaction model. There are, however, two differences between the test proposed by [26] and estimating (28).

First, the proposed test replaces the adult environment with a developmental input *i*_0_ = *−e*_0_ (which could be some feature of either the organism’s somatic state or its environment):

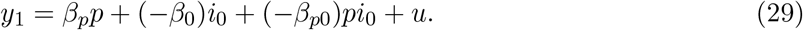

Second, [26] recommends implementing this analysis via a regression and visualization similar to the visualization that [14] suggests for the environmental mismatch hypothesis. Figure 4 provides an example of the sort of visualization [26] that might complement the regression in (29). For convenience, we shall call this empirical strategy a “developmental input-phenotype interaction approach.”

**Figure 4:**
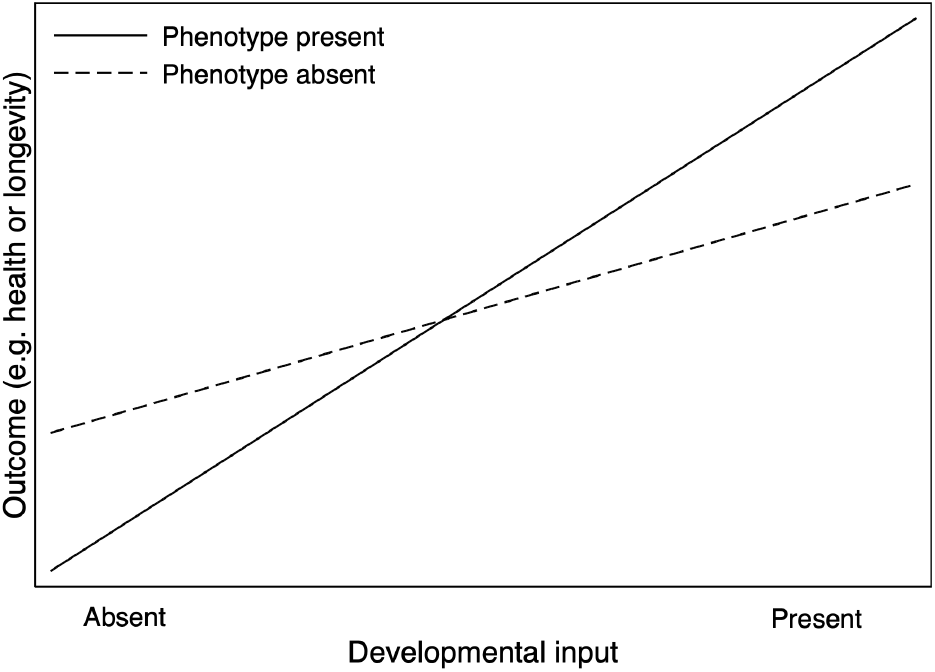
Visual depiction of an environment/phenotype interaction. This is conceptually similar to Figure 2 in the main text, but uses a developmental input on the x axis in place of adult environment. Figure adapted from [26].

We argue that the approach suggested in [26] has several problems. First, the visualization that motivates the interaction regression above suffers from a problem similar to the one in Figure 2 in the main text (a visualization of a developmental environment/adult environment interaction). Both versions flatten a three (or higher) dimensional problem into two dimensions. In doing so, the figures omit an important variable and thus force what are typically un-articulated and/or untested assumptions. Figure 2 omits the developmental environment because it conflates the potential effects of the developmental environment with the potential effects of a change in the environment. Figure 4 groups organisms for whom the developmental input is a correct prediction of adult state with those for whom it is not. The prediction depicted in the figure is correct for the former set of organisms, but not for the latter. The picture can still be valid when both sets of organisms are examined together, but only with the additional assumption that the developmental input is *on average* a good prediction of adult state.

If evolution is responsible for the studied form of predictive response/developmental plasticity, it is certainly true that the prediction must be correct more often than not, unless there was a recent change in the environment and selective landscape. If the predictions are often incorrect, then this form of plasticity should not evolve. However, building this assumption into the figure and resulting interpretation seems problematic for two reasons. One is that a central question in the study of developmental plasticity is the degree to which it is an evolutionary phenomenon [53]. The other is that the conflation of organisms for whom the developmental input is a good prediction and those for whom it is not reduces the sensitivity of the suggested test. A more precise figure would condition on the set of organisms it is addressing, and allow for the fact that the crossing property is mitigated (or may disappear) for the organisms with “mismatched” developmental input and adult somatic state.

Second, an interaction regression that uses adulthood instead of a developmental input might still suffer from omitted variable bias. This means that the combination of a test for a significant interaction between adaptation and adult somatic state/environment (*β*_*p*0_ ≠ 1) and the visualization will not yield an accurate test, because the estimate of *β*_*p*0_ will be biased.

Third, the interaction regression in (29) suffers the same potential omitted variable bias problem as the interaction regression (11) in the main text. (29) above takes a strong stance on the functional form of the relationship between (a) the developmental input and phenotypic adaptation and (b) the adult outcome. A safer assumption would be to start with a general functional form, e.g., *y* = *f* (*e*_0_, *p*^*a*^) and then take a Taylor approximation. In this view, a quadratic regression of the form

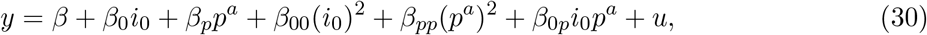

is safer. If this functional form is correct, then the interaction regression suffers omitted variable bias due to misspecification. Even if the interaction regression is specified correctly, then a quadratic regression would yield unbiased estimates of *β*_0*p*_, the trigger for a visualization. The test for whether the interaction regression was an acceptable specification is that estimates of *β*_00_ and *β*_*pp*_ are zero.

Fourth, because phenotypic adaptations are more likely be endogenous to an organism than adult environment is, the visualization and any regression that examines phenotypic adaptation rather than adult environment will be more likely to suffer selection bias (formally, a correlation between the explanatory variable–in this case, the phenotype–and the error term in the regression model). Consider an example in which the phenotype is how early an organism starts reproduction.

Both the visualization and interaction regression suggest that a test for internal PARs is whether those who receive a negative input during development start reproduction earlier than those who receive a positive input. This test requires that there be some individuals who receive the negative input who do start reproducing early, and some who do not; without such variation one cannot estimate the interaction term. But one has to have a theory for why two organisms with the same input have different responses. Unless that response is unrelated to any observable variables, e.g., is random, the coefficient on the phenotypic adaptation and the interaction term will suffer from selection bias. If there is selection bias, then the test proposed in [15] may cause one to reject the internal PAR hypothesis even if it is true.

Fifth, if DC is also true, but has a differential effect on organisms that undertake a phenotypic adaptation and those that do not (which is certainly a biologically plausible scenario), then the developmental input-phenotype interaction approach will be inaccurate. DC changes the slope of the adult outcome-developmental input line. If it operates more on organisms that undertake a phenotypic adaptation, then it may suppress cross-over suggestive of a PAR even when there is a PAR. If DC operates more on organisms that do *not* undertake the adaptation, then it may generate cross-over suggestive of a PAR even when there is no PAR.

Finally, we want to address an important point about any test that relies on a specific phenotype, rather than the mismatch test strategy, regardless of whether the test is theoretically consistent with a PAR definition. A phenotypic test takes a narrow view of phenotypic adaptations, with the consequence of reducing the test’s power to detect a PAR. Suppose that organisms who receive a negative developmental input can make several alternative adaptations. If the adaptations are substitutes for one another or mutually exclusive (due to, e.g., energetic or morphological constraints), those who adopt one adaptation will not adopt another and vice versa. If one only tests for one particular adaptation among those who receive the input, then one may reject PAR even when it is valid. In effect, if organisms that receive a developmental input can adopt a substitute phenotypic adaptation (*P* ^*′*^) than the one examined in Figure 4, and that alternative adaptation is as effective as *P*, then fitness without *P* will be the same as with *P*. This will also manifest in a non-significant interaction. In short, any phenotypic test that attaches itself to a specific adaptation carries the implicit assumption that there are not other, alternative adaptations.

### A.4 Conceptual issues with testing models

In the main text, we discuss two important conceptual issues that constrain tests of the DC, PAR, and AEQ hypotheses. These are the non-independence of the variables that are key to each theory 3.1, and the fact that the theories, as we have defined them, are not mutually exclusive 3.1.1. In this section, we discuss two others.

#### A.4.1 Overlapping predictions

A difficulty of testing the DC, PAR, and AEQ hypotheses is that they only generate divergent predictions under specific circumstances (Table 4). Even if we only examine two of the three hypotheses, only certain types of variation in developmental and adult environments allow us to distinguish one hypothesis from the other (Table 4). Specifically, PAR is not testable without variation in both the developmental *and* the adult environment [11, 38, 49]. There is no test that can distinguish PAR from DC unless there are subjects in both matching and non-matching conditions.

**Table 4:** Predicted relative health or fitness outcomes for an organism experiencing various early-life and adult environmental quality combinations, depending on whether the outcomes are generated by developmental constraints (DC), predictive adaptive responses (PARs), or both DC and PARs acting together.

|  | Environment | Only DC<br>is true | Only PAR<br>is true | Both DC &<br>PAR are true | Implication |
| --- | --- | --- | --- | --- | --- |
| (1) | HL vs HH | HL = HH | HL < HH | HL < HH | If $HL \geq HH$, can reject PAR & both |
| (2) | HL vs LL | HL > LL | HL < LL | Ambiguous | If $HL > (<) LL$, DC more (less) powerful than PAR |
| (3) | HL vs LH | HL > LH | HL = LH | HL > LH | If $HL \leq LH$, can reject DC & both |
| (4) | HH vs LL | HH > LL | HH = LL | HH > LL | If $HH \leq LL$, can reject DC & both |
| (5) | HH vs LH | HH > LH | HH > LH | HH > LH | If $HH \leq LH$, can reject all theories |
| (6) | LL vs LH | LL = LH | LL > LH | LL > LH | If $LL \leq LH$, can reject PAR & both |

H = high-quality environment, L = low-quality environment. The first letter in each pair is an organism’s developmental environment, while the second is their adult environment. E.g., row one compares health/fitness outcomes in individuals who had high-quality developmental but low-quality adult environments (HL) to those who had high-quality environments their whole lives (HH) (first column). The other columns explicate the predictions about individual health/fitness that follow if only DC were true (second column), only PARs were true (third column), or both were true (fourth column). < and > signs refer to which organism would have a better or worse health/fitness outcome in adulthood under the given hypothesis, respectively.

Relatively few observational studies can meet this challenging requirement, especially studies of human subjects. For example, the canonical studies of the Dutch Hunger Winter [54] and Leningrad Siege [55] only contain information on rows 5 and 2 of Table 4, respectively (though the degree to which adult environments matched developmental environments in Leningrad is open to interpretation [17]). This means that fewer human data sets than animal data sets are suitable for distinguishing PAR from DC because it is unusual to have access to human subjects in all the necessary developmental and adult conditions [but see 38, 56, 57, 58, 30, for exceptions].

#### A.4.2 Unobservable counterfactuals

While DC, PAR, and AEQ theories pertain to intra-individual differences in organismal responses to early and adult environments (see Section 2 in the main text), the reality is that tests of the theories will usually be executed using data on inter-individual differences in environment and outcomes. The same organism cannot, in general, be observed under two different developmental environments (*e*_0_), or in one situation where there are differences between their developmental and adult environments and another where there are not (∆*e*).

As a result, researchers must assume that environments are distributed randomly across subjects. In some situations this assumption is likely warranted, and in others not. For example, it is probably reasonable to assume that baboons who experience droughts are, on average, similar. However, it is less clear that this assumption is warranted for, e.g., human warfare; differences in resource access and social class strongly affect who bears the burden of such events. This issue is not specific to the theories in this paper, so we flag the problem but do not tackle it here. In the main text, our recommendations for testing the hypotheses assumes that researchers have access to data where environment is believed to be randomly distributed across organisms in the sample.

### A.5 Conditions under which it is possible to simultaneously test the developmental constraints (DC), predictive adaptive response (PAR), and adult environmental quality (AEQ) hypotheses

In Section 3.1 in the main text, we explain why it is generally impossible to test all three hypotheses (DC, PAR, and AEQ) simultaneously. It is impossible to manipulate subjects’ developmental environments, adult environments and the change in these environments independently; change in any one of these necessarily changes one of the others. However, there are two ways to redefine PAR to avoid this conceptual problem.

First, if one is specifically testing “symmetric” PAR rather than possibly “asymmetric” PAR, it may be possible to test PAR at same time as DC and AEQ. In symmetric PAR, positive and negative differences between developmental and adult environments both have the same magnitude and direction of effects on adult outcomes (see the left panel of Figure 3); in asymmetric PAR, one direction of environmental change (i.e., either positive or negative) more strongly influences adult outcomes than the other (see, e.g., the right panel of Figure 3). If we assume that PARs are symmetric and that the AEQ hypothesis is false, the data should show that positive and negative changes in environment each have identical negative effects on adult outcomes, as described in (19) and the left panel of Figure 3. However, if the AEQ hypothesis is true, then—even in the presence of symmetric PARs—the data should show that positive changes in the adult compared to the developmental environment have less negative effects on adult outcomes than negative changes in environment do:

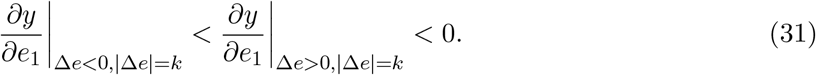

Thus, if we assume that symmetric PAR is true, then evidence of (31) is evidence for the AEQ hypothesis. However, if we are not *a priori* sure that PAR is symmetric, then evidence of (31) could be consistent with either (a) symmetric PAR plus AEQ or (b) asymmetric PAR where positive changes in the environment have less negative effects on adult outcomes than negative changes do (“left asymmetric PAR” as in right panel of Figure 3) and no AEQ. Indeed, one cannot even rule out the possibility of mild left asymmetric PAR that is made to appear *more* asymmetric by AEQ.

Building on this logic, it should also be clear that data ostensibly consistent with (31) (left panel of Figure 3) are also hard to interpret. If it is assumed that PARs are symmetric, then this can be interpreted as evidence against AEQ. But if it is assumed that PARs are right asymmetric, i.e., positive changes in environment are more harmful than negative changes in environment (e.g., the mirror image of the right panel of Figure 3), then this is evidence for the AEQ hypothesis. If one is not sure a priori whether PARs have symmetric effects or not, then one cannot rule in or out the AEQ hypothesis.

Second, if one alters the definition of PAR to focus on the difference between ancestral environment and adult environment rather than the difference between early-life and adult environment, it is possible simultaneously to test for PAR, DC and AEQ. The definition of PAR we employ in the main text has a developmental connotation that directly connects to DC, because both those depend on early-life environment, *e*_0_ (Table 1 in the main text). By contrast, an evolutionary PAR hypothesis may be defined as

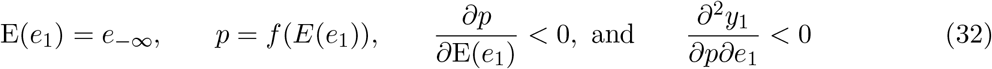

This definition of evolutionary PAR is similar to the definition of developmental PAR in Table 1 in the main text, except that the organism’s expectation of adult environment depends on ancestral environment (*e*_∞_) rather than its own early-life environment (*e*_0_). This change implies that, for evolutionary PAR, the phenotypic test is *∂p/∂e*_−∞_ *<* 0 and the mismatch test is *∂y*_1_*/∂*|*e*_1_ *− e*_−∞_| *<* 0. Because these tests do not depend on *e*_0_, it is possible to manipulate simultaneously *e*_0_ to test DC, *e*_1_ to test AEQ, and, e.g., |*e*_1_ *− e*_−∞_| to test evolutionary PAR. Changes in *e*_0_ and *e*_1_ do not dictate changes in |*e*_1_ *− e*_−∞_|, so there is no identification problem.

Luke and colleagues [35] provide an example of a mismatch test for evolutionary PAR that illustrates this last point. They regress adult body mass index (*y*_1_) on ancestral (*e*_−∞_) and adult incomes (*e*_1_), with local crop productivity serving as a proxy for ancestral income. This specification implicitly tests for evolutionary PAR and AEQ. The analysis could also have tested for DC by adding childhood (or parental) income because ancestral income has variation (the difference between potential income implied by ancestral income and the actual early-life income) that is independent of early-life income.

This second change in the definition of PAR does not, however, disprove our basic claim, that one cannot simultaneously test developmental PAR, DC and AEQ. To put a finer point on it, if one also changes the definition of DC to focus on ancestral environment, i.e., define an evolutionary DC hypothesis as a negative effect of bad ancestral environment or *∂y*_1_*/∂e*_−∞_ *>* 0, then one will not be able simultaneously to test for evolutionary DC, evolutionary PAR, and AEQ.

### A.6 Additional notes on the conceptual problems with the interaction visualization

In Section 4 in the main text, we discuss problems with separately plotting adult outcomes on adult environment for individuals who started in high-quality and low-quality developmental environments (Figure 2 in the main text). Other papers have noted the conceptual difficulty with testing for PARs when researchers only have data on subjects that all experienced similar adult environments, as is the case in some canonical human studies [e.g., 59]. These papers note that it is impossible to know whether individuals who started in high-quality environments end up doing better because their developmental environment was high-quality, or because their developmental and adult environments match [33, 11, 49, 38]. While this is not strictly a criticism of plots like Figure 2 in the main text, it can be interpreted as an argument that Figure 2 cannot be plotted if there is no variation in adult environment.

However, if a visualization like Figure 2 can be plotted, i.e., subjects experienced variation in their adult environments, then one has to address whether Figure 2 and depictions like it are informative. Plotting outcomes against adult environments is a cumbersome way to visualize PARs. It does not cleanly isolate the effects of starting point (i.e., developmental constraints), because it uses the variable of interest from yet a third hypothesis (the adult environmental quality hypothesis) as the x axis. Moreover, there is ambiguity about what pattern should be interpreted as evidence for PARs. The bisection from above of (a) the adult outcome-adult environment line for organisms from high-quality developmental environments by (b) the same line for organisms from low-quality developmental environments, such as illustrated in Figure 2, is a sufficient but not necessary condition for PARs. A sufficient condition is that the slope of the low-quality line is smaller than the slope of the high-quality line. Finally, if a plot like Figure 2 is generated with predictions from estimation of an interaction model (as opposed to plots of raw data), either the plot cannot find PARs or suffers from omitted variable bias and may not be informative. The reasons for this are discussed in detail in Section 6 in the main text.

### A.7 Additional implications for the interaction model

Here we discuss two additional problems with using the interaction regression (Eq. 11) to test for PARs and DC.

One problem is that the appropriate test for DC when using a univariate regression of outcome on developmental environment is not the correct test for DC in the interaction regression model. A reasonable test for DC in a regression where adult outcome is the dependent variable is whether the coefficient on the predictor variable *e*_0_ is positive. That is an appropriate test if one estimates a model along the lines of

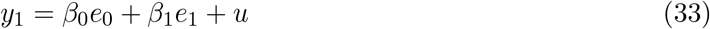

(assuming that such a model is the correct specification). Recall that the definition of DC implies that the test for DC is that derivative *∂y*_1_*/∂e*_0_ is positive. In the model above (Eq. 33), the derivative equals *β*_0_. But if we are estimating an interaction regression as in equation 11, the correct derivative is instead *β*_0_ + *β*_01_*e*_1_. Even if *β*_0_ *>* 0, the derivative could be negative if *β*_01_*e*_1_ *> −β*_0_.

Another problem is that if the interaction regression is not a correct description of reality (e.g., if the quadratic regression is actually correct), then the coefficients estimated by the interaction regression suffer from omitted variable bias. Compared to the quadratic model, the interaction regression omits squared 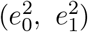 and interaction (*e*_0_|∆*e*|) terms. With ordinary least squares estimation, other coefficients to some extent absorb variance that should be attributed to omitted terms. As a result, the coefficient on the interaction is biased:

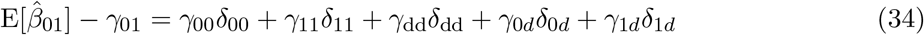

where *δ*_xz_, for *x, z* ∈ {0, 1, *d*}, is the coefficient on *e*_*x*_*e*_*y*_ from a regression of *e*_0_*e*_1_ on *e*_0_, *e*_1_, and *e*_*x*_*e*_*z*_. If one were to test whether PARs should be rejected by testing whether *β*_01_ = 0, the test may be inaccurate. The test could reject that *β*_01_ = 0 even though *γ*_01_, the true measure of the interaction, is zero. Likewise, it could fail to reject *β*_01_ = 0 even though *γ*_01_ is zero. When testing for DC, the coefficient on both the interaction and the *e*_0_ term may be biased for similar reasons, leading to potentially incorrect results even if the test for DC (Eq. 1) is employed. Even if the test obtains the right answer, it produces inaccurate conclusions about effect sizes.

The interaction regression would not produce omitted variable bias if the two variables in the interaction term were both truly binary. In that case, the squared terms are identical to, and thus fully captured by, the non-squared terms. However, the environment is rarely binary. For example, a variable such as drought/no drought is a continuous rainfall variable that has been transformed into a categorical variable. This is true even if environment is a continuous variable that one cannot measure continuously and therefore treats as categorical. An example is the Dutch Hunger Winter: we do not have data on actual caloric consumption during the Winter, but we know that it was lower than during the winters before or after it. Whether the continuous variation is measurable or not, a categorical partitioning measures the underlying continuous variable with error. Even if the interaction regression were a true model of the world, the coefficients on the categorical environmental variables would estimate the effect of changes in environment with bias from the measurement error.

The implication of these biases is that, if the interaction model is used to test these theories, it is likely to produce high rates of false positives and false negatives. That is, it is likely to cause us to conclude that PAR and/or DC are true when they are not, and/or that they are not true when they actually are. We explore this via simulation in the next section.

### A.8 Baboon data

In order to ensure that we selected covariate values that are representative of a real-world biological system, we summarized long-term data from the Amboseli Baboon Research Project and used the resulting ∆*e* values in our simulations (i.e., the value of the difference between developmental and adult environments). The subjects were wild female savannah baboons (primarily *Papio cynocephalus* with some naturally occurring admixture from neighboring *Papio anubis* populations) living in the Amboseli ecosystem in southern Kenya. This baboon population has been studied on a near-daily basis since 1971 by the ABRP, which collects longitudinal demographic, ecological, life history, and behavioral data on individually known animals [42]. The data we summarized to obtain values used in the simulations spanned January 1980 to March 2021.

Like many other species including humans, baboons can experience a variety of developmental environments [33, 60, 61, 32]. We summarized a major source of variation in the developmental environment for baboons: dominance rank. Female baboons form strong, stable dominance hierarchies that determine access to important resources [62]. Female infants “inherit” their ranks from their mothers via a system of youngest ascendancy, where the infant becomes dominant over any older maternal sisters at birth. The functional consequence of the youngest ascendancy system is that female baboons can reasonably predict that they will hold a dominance rank similar to the one their mother held when they were born, when they themselves are adults. However, due to events such as group fissions and matriline overthrows [42], some animals occupy dominance ranks in adulthood that are not similar to the dominance rank their mother held when they were born. Since there is variation in both developmental environment (some baboons are born to low-ranking mothers, and some to high-ranking ones) and the how well the developmental and adult environments match (some females will hold nearly the same rank their whole lives, and some will experience change), rank would be an appropriate variable for testing the DC and PAR hypotheses. We used relative (i.e., proportional) dominance ranks, which represent the proportion of adult female group members that the subject in question outranks [63, 62]. For example, a dominance rank of 0.9 means that the female outranks 90% of the adult females in her group. Each animal’s rank was determined based on the outcomes of all observed, decided agonistic interactions between adult females. Trained observers, who could recognize individual animals via differences in morphological characteristics, recorded the identities of individuals participating in agonistic encounters and the outcome of each of these encounters. If one animal behaved submissively while the other was either aggressive or remained neutral, the interaction was recorded as “decided” with respect to the rank relationship between the two interacting animals. Any interactions without a clear outcome (e.g., where both animals displayed submissive signals) were excluded from dominance rank calculations.

For rank during development, each female subject was assigned the rank her mother held in the month the subject was born. In order to summarize the difference between developmental and adult rank, we needed time window(s) during adulthood in which to capture rank. We eliminated periods of rank instability (e.g., when groups were going through fissions, during which rank can be difficult to determine), then aggregated information about months in which females were cycling (i.e., months in which females were not either pregnant or nursing). A data set like this would be suitable for testing if developmental environments, or the difference between developmental and adult environments, predicted (for example) the chances that a female would successfully conceive, or the number of mating consortships she participated in. The difference between a female’s mother’s rank the month she was born and the rank she held during the cycling month(s) in question was the difference between her developmental and adult environment (*δe*). This resulted in a data set that contained information about 7,661 months for 281 individual female baboons. After weighting each baboon equally, the mean difference between developmental and adult rank was −0.03 (SD=0.21, range=-0.95-1). Data sets constructed across the course of pregnancies or lactation resulted in very similar values (e.g. for pregnancies mean =-0.02, SD=0.22, range=-0.92-1). We chose to use the summarized values obtained from cycling months since it was the largest of the data sets.

### A.9 Further details on the simulations

This section provides more details on 4 of the 5 steps in our simulations.

#### 1. Generating different virtual realities

We prepared a set of possible “realities” where the PAR hypothesis was either true or false. We started with the 3rd-order polynomial for *y*_1_ = *f* (*e*_0_, |∆*e*|), which generated an equation with 10 coefficients:

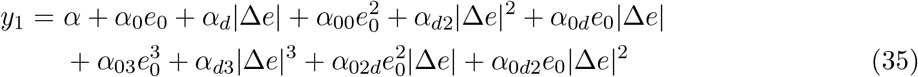

Each reality is defined by (a) a 10 × 1 vector *θ ∈* Θ = {*α, α*_0_, *α*_*d*_, *α*_02_, *α*_*d*2_, *α*_0*d*_, *α*_03_, *α*_*d*3_, *α*_02*d*_, *α*_0*d*2_} combined with (b) the equation above that relates early environments and change in environment to adult outcomes.

Why do we assume reality is described by a polynomial and why specifically a third-order polynomial? Reality permits a general function relating *y*_1_ to *e*_0_ and |∆*e*|: *y* = *f* (*e*_0_, *e*_1_, |∆*e*—). To make realistic virtual realities that are tractable for simulation, we took a Taylor series expansion of that general function around zero. We needed to select a finite order at which to stop the expansion. A 1st-order polynomial would cause the interaction term in an interaction regression to always be zero. A second-order term would always be estimated well by a quadratic and not an interaction, but by assumption (i.e., we would be simulating a world in which our suggested empirical test, by definition, would always perform perfectly). A 3rd- or higher-order polynomial allows both the interaction and quadratic to generate errors that can be compared, which is our goal. However, as polynomial order increases, exponentially more computation power is required to simulate reality given a fixed level of variation in (*e*_0_, |∆*e*|). ^8^ A 3rd order polynomial is a compromise between a model that is more general and one that is computationally tractable.

Even a 3rd-order polynomial can have infinitely different coefficient values and thus describe infinitely different realities. We pared those realities down by, first, only simulating realities on the nodes of a 10-dimensional lattice in Θ where the value of each dimension takes 5 values ranging from −1 to 1. Specifically, we created a grid of points in a 10-dimensional space. The points in each dimension took value (−1,−0.5,0,0.5,1), so the grid had 5^10^ points. This grid represents possible realities defined by different values of coefficients on up to 3 powers of (*e*_0_, |∆*e*|), i.e., each reality is a 10 × 1 vector *θ*. We restricted the range of coefficients to [*−*1, 1] because these are consistent with either binary outcomes or continuous, bounded outcomes, as well as environmental variables that range from 0 to 1.^9^ We chose 5 values in the [-1,1] range so that the grid was not so dense that there were too many possible realities to simulate.

Second, we ruled out realities where at any value of *e*_0_ *∈* [0, 1] or ∆*e ∈* [*−*1, 1] generated an outcome outside the range of [0, 1]. That is, we retained outcomes that could map onto binary outcomes (0/1), or onto outcomes which are continuous but bounded.^10^ To implement this, we allowed **x** = (*e*_0_, |∆*e*|) to each take 4 possible values (0, 1/3, 2/3, 1), and calculated the 16 possible outcomes that resulted from each coefficient vector *θ*. If any outcomes were outside [0, 1], we rejected that coefficient set. This filtering produced 130,201 parameter combinations and thus “pruned” realities.

#### 2. Determining whether PAR or DC are true in each reality

Each of these feasible realities was evaluated for PARs and DC by applying the tests in equations 7 and 1, respectively. With a 3rd-order polynomial, the derivatives in these tests depend on the value of *e*_0_ and ∆*e*. We evaluated the derivatives for each reality at 16 evenly spaced values of (*e*_0_, ∆*e*), by picking four evenly spaced values for *e*_0_ and for *e*_1_, and calculated ∆*e* from these. A reality was labelled positive for PARs or DC if the relevant derivative test was satisfied on average across these values. Of the 130,201 pruned realities, 2,697 (2.07%) were truly positive for PARs, 2,697 (2.07%) were truly positive for DC, and 58 (0.04%) were truly positive for both.

#### 3. Simulating a data set for each reality

We created a simulated data set for each *θ ∈* Θ^*f*^ that corresponded to a pruned reality. Each simulated data contained a set of 2000 observations on (*ŷ*_1_, *e*_0_, ∆*e*). We start by explaining how we generated the environmental variables in the simulated data. Then, we explain how we generated the observed outcome variable *ŷ*_1_.

Values of early environment and change in environment were selected to reflect patterns from the real world, namely social status of females (i.e., dominance rank) in a natural population of wild baboons in Kenya that are the subject of long-term, individual-based monitoring by the Amboseli Baboon Research Project in Kenya [42]. While social status predicts some outcomes for females (e.g., access to resources and some fertility measures [62]), our goal was not to conduct a full empirical test of the DC and PAR hypotheses using these data. Rather, we wanted to use these values to provide a realistic range of developmental and adult environments for the simulations. Usefully, the observed distribution of developmental dominance rank (*e*_0_) and the difference between subjects’ developmental and adult ranks (∆*e*) generate covariate values that are highly generalizable to many kinds of environmental quality measures in a wide variety of species. In our data, *e*_0_ had a roughly even distribution spanning the range from 0 (lowest-rank or worst) to 1 (highest-rank or best), and ∆*e* had an approximately normal distribution in the range [*−*1, 1] with a mean of *−*0.04 and a standard deviation of 0.21. More details about these data can be found in the supplement (A.8).

With this as background, we generated our environmental variables in two steps. First, we drew 100 evenly spaced values of *e*_0_ *∈* [0, 1]. Second, for each *e*_0_ we drew an associated 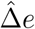 from a normal distribution with mean 0.04 and standard deviation 0.21. Third, we took 3 steps to ensure that the implied *e*_1_ fell in the range [0, 1]. We computed 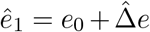 for each *e*_0_; computed a new, evenly-spaced *e*_1_ based on rank order of *ê*_1_; and then computed ∆*e* = *e*_1_ *− e*_0_. The result was 100 different realistic and feasible combinations of (*e*_0_, ∆*e*).

Next, we generated we generated 20 values of observed outcome *ŷ*_1_ for each combination of (*e*_0_, ∆*e*). To do this we obtain the true adult outcome *y*_1_ implied by a given (*e*_0_, ∆*e*) and the pruned reality. Then we take 20 draws on an error *v* from a normal distribution with mean 0 and variance equal to *A*(1 *− A*), where *A* is the true outcome *y*_1_ at *e*_0_ = 0.5 and ∆*e* = 0 in a pruned reality. The variance is modeled on the variance of a binomial variable (like fertility) with mean *A* and *A* is modeled on the outcome of the mean female in a baboon group. We finish this step by generating 20 values of observed *ŷ*_1_ using the equation

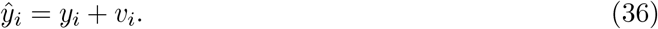

The result is a data set with 2000 observations: 20 observations on *ŷ*_1_ for each of 100 combinations of (*e*_0_, ∆*e*).

### Testing for DC and PAR in each reality

We estimated an interaction regression and quadratic regression on the N=2000 simulated data set for each reality. From the regression results for each reality, we generated four test results for PARs and three test results for DC in that reality (described in Table 2 and Section 7.1 in the main text). We then compared these test results to the truth for each reality (determined in the prior step) to estimate the sensitivity and specificity of each test for PAR and DC. We also tested for PARs and DC in each reality using a coin flip, simulated with a Bernoulli random variable with mean 0.5. The comparison allowed us to calculate the fraction of realities where each of the four tests, along with the coin flip, generated results that differed from the ground truth (Table 2 in the main text).

The simulation code is available online at https://github.com/anup-malani/PAR/blob/d3d9588af6bb96c49f0550c94aa756e6a1261a9f/PAR_simulation_220117b.do.

## Footnotes

1 The developmental mismatch hypothesis should not be confused with the evolutionary mismatch hypothesis [31], which replaces early-life environment with environment during some historical time period and stresses the mismatch between ancestral environment and current environment. We discuss this version in the supplement (A.5).

2 A technical caveat: opposite sign changes can decrease the mismatch if one starts from where *e*_0_ *> e*_1_. Therefore, the mismatch hypothesis may require the assumption that *e*_0_ *≤ e*_1_ to be a valid test of PAR. When that is not true, mismatch may be true but needs a theoretical justification aside from PAR. This is related to the limited environments in which one can test for PAR listed in Table 4.

3 In the supplement (Section A.5) we give more details about the conditions under which all three hypotheses are testable. One is the exception given in the main text. The other is when one defines an evolutionary version of PAR focused on the differences between ancestral and current environments [35].

4 If the low-quality developmental environment line merely has a lower slope than the high-quality developmental line, the plot can be interpreted as evidence for a PAR: the lines do not need to cross, and may be likely not to if developmental constraints are also at work [e.g., 14].

5 Values are motivated by the mean and SD from data on a natural population of wild baboons in Kenya that are the subject of long-term, individual-based monitoring by the Amboseli Baboon Research Project in Kenya [42]. Details can be found in Section A.8.

6 The term immediate adaptive response is in use elsewhere in the literature [e.g., 18, 46]. It is sometimes used as a synonym for developmental constraints, so we deliberately avoid using it here.

7 It is true that a developmental adaptive response (DAR) could trigger adaptation. However, the problem there is that PAR and DAR are indistinguishable, not that the test for PAR is incorrect. Another way to put this is that the test for *β <* 0 is a test for both PAR and DAR, not that omitted variable bias means that this test is inaccurate. This is discussed in detail in Section A.2.

8 For example, a 4th order polynomial has 15 terms; because we consider 5 values of each coefficient in the polynomial, a 4th order polynomial requires consideration of 5^15^ rather than 5^10^ realities.

9 Note that the range of coefficients is not the same as the range of, e.g., *e*_0_. Instead it is the range of possible causal impact of powers of environmental quality on outcomes.

10 An affine function of a bounded, continuous variable can transform it into continuous variable with a range of [0,1]. Moreover, the test for PARs or DC can be adjusted to account for that transformation.

